# Modeling recent positive selection in Americans of European ancestry

**DOI:** 10.1101/2023.11.13.566947

**Authors:** Seth D. Temple, Ryan K. Waples, Sharon R. Browning

## Abstract

Recent positive selection can result in an excess of long identity-by-descent (IBD) haplotype segments. The statistical methods that we propose here address three major objectives in studying selective sweeps: scanning for regions of interest, identifying possible sweeping alleles, and estimating a selection coefficient *s*. First, we implement a selection scan to locate regions of excess IBD rate. Second, we develop a statistic to rank alleles that are in strong linkage disequilibrium with a putative sweeping allele. We aggregate these scores to estimate the allele frequency of the sweeping allele, even if it is not genotyped. Third, we propose an estimator for the selection coefficient and quantify uncertainty using the parametric bootstrap. Comparing against state-of-the-art methods in extensive simulations, we show that our methods are better at identifying sweeping alleles that are at low frequency and at estimating *S* when *S* ≥ 0.015. We apply these methods to study positive selection in European ancestry samples from the TOPMed project. We analyze eight loci where the IBD rate is more than four standard deviations above the population median. The IBD rate at LCT is thirty-five standard deviations above the population median, and our estimates of its selection coefficient imply strong selection within the past two hundred generations. Overall, we present robust and accurate approaches to study very recent adaptive evolution without knowing the identity of the causal allele or using time series data.

## Introduction

Positive natural selection favors alleles advantageous for survival or reproduction. The process of evolution via positive selection is often modeled via hard or soft selective sweeps. In hard sweeps, a new mutation increases rapidly in frequency because it provides a significant evolutionary benefit. In soft sweeps, an existing neutral allele at low frequency increases in frequency when it becomes newly beneficial in the context of environmental change^1–3^. The sweep is partial or incomplete if the selected allele is not fixed in the present-day. Alleles on the same haplotype as a sweeping allele increase in frequency as well until crossover recombination separates them from the sweeping allele. Motivated by this observation of genetic hitchhiking, most methods to detect and model recent positive selection evaluate patterns of unusual linkage disequilibrium^4^.

Selective sweep events in the past five hundred generations can leave profound footprints on genomic diversity around a selected locus. For instance, extended haplotype homozygosity about the LCT gene in northern Europeans spans multiple megabases (Mb) of genetic sequence^5^. Studying positive selection can be complementary to large association studies; for instance, loci associated with inflammatory disorders show evidence of strong and weak selection^6^. The purpose, magnitude, and prevalence of such adaptive evolution remain open questions in population genetics, medical genetics, and conservation genetics.

The popular mathematical model to study a selective sweep from a single allele frames the problem in terms of a selection coefficient *s*. This parameter measures the relative advantage that the sweeping allele has over alternative alleles. In the haploid model, the deterministic formula *p*_*s*_(*t* − 1) = *p*_*s*_(*t*) × (1 + *s*)/(1 + *p*_*s*_(*t*) × *s*) describes how the allele frequency changes over a single generation from time *t* to t-1 generations ago. Similar formulas for diploids assuming additive, multiplicative, dominance, and recessive genetic models are in Appendix A. These functions may be inverted to express the allele frequency *p*_*s*_(*t*) one generation further back in time in terms of the allele frequency *p*_*s*_(*t* − 1) and the selection coefficient *s*. Together selection and genetic drift imply a probability distribution on the historical frequencies of the sweeping allele.

To scan for targets of recent positive selection, standard methods study either decay patterns of extended haplotypes^5,7^, excess recent shared ancestry^8–11^, changes to the site frequency spectrum^12^, haplotype differences between ancestrally diverged populations^13,14^, allele frequencies in ancient DNA samples^15^, or excess ancestry proportions in admixed samples^16^.

Here we take an identity-by-descent (IBD) based approach. Detection of IBD segments can be accurate if the shared segment spans a length of two or more centiMorgans (cM)^8,9,17–21^. We define “recent” to mean selection within the last five hundred generations, the time period in which such long IBD segments derive from their common ancestor. Our methods examine very recent partial sweeps; haplotype homozygosity methods study slightly older selection within a population; differences between populations relate to even older selection.

To identify possible sweeping alleles, the earliest method proposed is extended haplotype homozygosity (EHH). This statistic measures the proportion of shared haplotypes at increasing physical distances^5^. Additional methods derive from this initial idea^7,14,22–24^. All these approaches compute statistics derived from population genetics theory, normalize over the genome, catalog regions where these statistics differ from a genome-wide central tendency, and rank alleles as good candidates for selection based on the magnitude of these deviations. They do not estimate the frequency of the unknown sweeping allele.

To estimate the selection coefficient, four different paradigms have been considered. Mathieson and McVean (2013)^25^ and Mathieson and Terhorst (2022)^26^ use changes in allele frequency over time to estimate a constant selection coefficient or time-varying selection coefficients. These methods are limited to studies where data is collected at multiple time points. The coalescent defines a model for how samples relate to each other in the context of an ancestral tree^27,28^, and coalescent-based approaches can be used to estimate the selection coefficient using either present-day data^29,30^ or time series data^31^. Coalescent-based approaches are sensitive to the initial task of ancestral tree estimation. Some tree inference algorithms apply ad-hoc rules to handle increasing sample sizes, but these methods do not quantify uncertainty over the space of possible tree sequences^32–34^. Alternative approaches express tree uncertainty by sampling from a Bayesian posterior distribution, but these approaches scale poorly with increasing biobank sizes^35–37^. Machine learning approaches train neural networks based on simulated genetic data for known selection coefficients as opposed to traditional statistical inference^38–40^. Predictive accuracy depends on the model architecture and training.

Another concern in evolutionary biology is that alternative mechanisms could explain the patterns of haplotype structure that these methods leverage. For instance, authors repeatedly suggest that the selection signal at the major histocompatibility complex (MHC) found in many European population studies is an example of balancing rather than positive selection ^6,41^. Further confounding factors include population structure, assortative mating, and unaccounted for familial relatedness. These issues can be partially addressed by preliminary data analysis^42–44^. Some existing methods require ancient or archaic samples that are available for Eurasian populations but limited for other populations or non-human studies. A complete methodology that is motivated by statistical and population genetics theory, makes as few assumptions as possible, and is easily interpretable would broadly contribute to the research community.

We propose a suite of methods to study all three aspects of a selective sweep analysis from a sample of whole genome sequences. This inference framework examines the effects of strong selection (*s* ≥ 0.015) on identity-by-descent within a population in the last five hundred generations. In a simulation study on IBD tracts drawn from the coalescent model, we evaluate the accuracy of selection coefficient estimation for varying allele frequencies, demographic scenarios, and model misspecifications. In a simulation study on IBD tracts inferred from sequence data, we compare our approach alongside state-of-the-art methods for identifying the sweeping allele and estimating its selection coefficient. We implement our estimation procedure to investigate positive selection in European Americans in the TOPMed project^45^. Annotations of the eight genes identified in our IBD-based selection scan highlight pigmentation and immune response as possible targets of selection in Europeans.

## Materials and Methods

### Overview of Methods

Here we offer a unifying framework for recent and strong selective sweeps that connects shared haplotypes to a statistical model for recent ancestry. At a locus, haplotypes carrying and not carrying the sweeping allele bifurcate into two coalescent subtrees^46,47^. The subtree with the adaptive allele coalesces more quickly than expected given genome-wide relatedness. This imbalance is more profound the greater the selection coefficient. IBD tracts overlapping a locus provide information about the underlying coalescent tree.

Selective sweeps can create extremely high local IBD rates and large clusters of IBD haplotypes compared to average genome-wide relatedness. We survey the genome for loci with an excess of inferred IBD tracts as in Browning and Browning (2020)^8^. At loci with elevated IBD rates, we infer a haplotype group of excess IBD, and then we calculate scores for alleles based on their frequency difference between the inferred group and the rest of the sample. These scores serve as the basis for ranking alleles as possible targets of selection. Using these scores, we produce weighted average estimates for the location and frequency of the sweeping allele.

Next, conditional on the allele frequency, we propose an estimator for the selection coefficient that relates the amount of IBD to the coalescent model under selection. Finally, we perform a fast parametric bootstrap procedure to calculate confidence intervals (CIs) for the selection coefficient. We implement this suite of methods iSWEEP (**i**ncomplete **S**elective sweep **W**ith **E**xtended haplotype **E**stimation **P**rocedure) in freely available software.

### Detecting IBD Segments

Our methods use IBD segments inferred from genetic data. We use two existing software programs to infer IBD tracts: hap-ibd^17^ and ibd-ends^8^. The accuracy of IBD segment detection depends on the cM length threshold: longer segments are more reliably inferred. We use different length thresholds in different contexts. We use segments longer than 2.0 cM to scan for selected loci and estimate the selection coefficient, and we use segments longer than 1.0 cM to infer an outgroup of excess IBD sharing. Estimating the selection coefficient is sensitive to inaccuracies in segment detection, whereas determining a haplotype cluster is an intermediate step in our procedure that tolerates more false positive segments. We specify algorithm settings for detecting IBD segments longer than 2.0 and 1.0 cM in Supplementary Table 1.

### Scanning for Excess IBD Rates

The number of IBD tracts longer than 2.0 cM divided by the number of haplotype pairs is the IBD rate we use in our scan. Every twenty kilobases (kb) we calculate the IBD rate based on detected segments that overlap the focal position. Positions where the IBD rate exceeds the genome-wide median plus four times the standard deviation are considered loci with excess IBD rate. Compared to the scan in Albrechtsen, Moltke, and Nielsen (2010)^11^, this scan concerns long IBD segments overlapping a focal position, rather than the probability that alleles are IBD at a marker.

Thus far our approach mirrors the selection scan in Browning and Browning (2020)^8^. We make three additional modifications. One change is that in calculating the genome-wide standard deviation we exclude positions that are initially three standard deviations below the median.

Another change is that we ignore positions within 0.5 cM of the start and end of the recombination map for each chromosome. These quality control measures exclude data near telomeres and centromeres where IBD calling is difficult. Lastly, we require that that a genomic region of interest span a 1.0 cM contiguous stretch of positions where the IBD segment counts exceed the threshold. Loci that only exceed our threshold for a few positions may be false positive results.

### Inferring an Outgroup of Excess IBD

Haplotypes descendant from an adaptive haplotype may belong to large clusters if selection is recent and strong. We develop a procedure to infer an outgroup, denoted *A*, with excess IBD sharing compared to the rest of the sample, denoted *B*. First, we detect IBD segments with length 1.0 cM or greater that overlap a focal location. Let haplotypes be nodes and IBD segments be edges in the context of a network graph. Because IBD segments longer than 1.0 cM are infrequent in a population sample, this graph should contain many disconnected haplotype clusters^48^. We also impose the rule that a node in a cluster may only be three edges away from its highest degree node (the haplotype sharing the greatest number of IBD segments). This condition ensures that there are no bridging edges on the periphery of clusters that would otherwise be disconnected.

Sizes of these haplotype clusters amount to an empirical distribution for IBD cluster size. We compute a sample mean and standard deviation for cluster size. Then, we identify clusters whose size exceeds the mean plus three times the standard deviation. These inferred clusters â together form the outgroup Â; the rest of the sample we denote as *B*^^^. We apply the IBD outgroup algorithm twice in our analysis, and we denote Â_1_, Â_2_ and *B*^^^_1_, *B*^^^_2_ for the first and second applications.

### Pinpointing the Focal Position of the Sweeping Allele

Genomic regions passing our selection scan encompass many Mb of genetic sequence. The IBD outgroup algorithm is sensitive to the initial location of IBD detection. We develop an intermediate step to focus on smaller regions where the sweeping allele may lie. Variants hitchhiking on the sweeping allele should segregate at similar frequency, suppressing common variation locally about the position of the causal allele. We take a sliding window approach to localize the signal. Consider *Q*_*w*_ = ∑^10^_k=1_ *p*_(*k*)_ where is *p*_(*k*)_ is the *k*^*t*ℎ^ smallest percentile of allele frequencies in the window *w*. Allele frequencies are not derived allele frequencies, but rather those of the allele more frequent in the inferred group Â_1_. High *Q*_*w*_ values correspond to windows where common variation is greater than that of a site frequency spectrum at neutrality.

The window with the highest *Q*_*w*_is a refined focal location for further steps in the estimation procedure. We use window sizes and step sizes of 250 kb and 50 kb, respectively. To address low marker density regions, we do not consider windows where there is a gap of 25 kb without SNPs.

We contrast our approach to that of the singleton density score from Field et al. (2016)^49^.

Their approach computes the physical distance between singleton variants. They note that the coalescent model with strong positive selection has long terminal branches, which should result in more *de novo* mutations close to each other. Singleton density score is based on rare variants; *Q*_*w*_ is based on common variants being at intermediate to high frequency in a window. We only use this statistic to refine a location for IBD calling, whereas singleton density score is applied to genome-wide selection scans.

### Estimating the Sweep Frequency and Location

At the refined location, we apply the IBD outgroup algorithm to infer the Â_2_ group with excess IBD sharing. For each variant *j* within 150 kb of the refined location, we compute a scaled difference in frequencies. Let *p*_*j*_ and *p*_*j*_,Â_2_ be the present-day allele frequency in the entire sample and in the inferred outgroup Â_2_, respectively. The scaled difference in frequencies is:

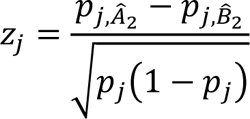

Except for a constant scalar, these scores represent the two-sample difference in proportions test statistic. They are similar to the change in derived allele frequency (ΔDAF) statistic in Grossman et al. (2010)^50^. However, the ΔDAF statistic is based on derived alleles between populations, whereas we consider a detected outgroup within a single population. We rank SNPs as candidates for selection (or in strong LD with a sweeping allele) based on the *Z*_*j*_ scores.

We aggregate evidence across variants to estimate a frequency for the selected allele. We take a sliding window approach in terms of physical distance and allele frequency. We consider rectangles of allele frequency 10% length and 500 kb width, and we move them by increments of 2.5% and 50 kb. For each rectangle, we average the *Z*_*j*_ variant scores contained in it. To address sparse marker coverage, we require that rectangles include at least five markers. The middle frequency and basepair of the rectangle with the highest average *Z*_*j*_ are our estimates of the selected allele’s frequency and location.

### Information Entropy Criterion Supporting the Sweep Model

Loci that pass our selection scan may not be the result of a selective sweep. Other biological processes may give rise to excess IBD rates. For instance, under balancing selection, there could be multiple beneficial haplotypes that each maintain appreciable frequency. In the recent and strong sweep model, we expect elevated extended haplotype homozygosity. Let *q*_â_ be the proportion of haplotypes in cluster â out of inferred outgroup Â. We propose to use the information entropy statistic ∑_â∈_Â−*q*_â_ log_10_ *q*_â_. The entropy increases with more clusters and/or decreasing size of the largest cluster. For example, if Â consists of one cluster, the entropy is 0.00; if Â consists of two equally sized clusters, the entropy is 0.30; and, if Â consists of three equally sized clusters, the entropy is 0.48. IBD entropy greater than 0.50 is supporting evidence for a sweeping allele that is at high frequency in the present-day.

### Estimation of Selection Coefficients

We estimate the selection coefficient using genetic data collected from the current generation. Let the IBD tract length *L* be a random variable that depends on a genetic model for selection and its coefficient *s*, the allele frequency *p*(0), and effective population sizes *N*_*e*_(*t*). The probability that the IBD tract length *L* exceeds a specified cM threshold is derived in Appendix A. This probability is identically distributed for each pairwise IBD tract in the population sample, and the expected count of such long IBD tracts is this probability times the sample size of pairs. Fixing *N*_*e*_(*t*) and *p*(0), we optimize for the selection coefficient *s*^ that minimizes the squared difference between the expected and observed number of IBD tracts longer than 3.0 cM.

To estimate a confidence interval for the selection coefficient, we simulate IBD tracts from a coalescent model parameterized in *s*^, *p*(0), *N*_*e*_(*t*), and the genetic model for selection. This simulation amounts to computing shared recombination endpoints that are drawn from recent common ancestors. Next, we re-estimate selection coefficients Ŝ_1:*B*_for *B* bootstraps. We compute the confidence interval Ŝ ± σ^^^_1:*B*_ × *q*_α/2_, where σ^^^_1:*B*_ is the sample standard deviation of bootstrap estimates and *q*_α/2_is the α/2 quantile of the standard normal distribution. An efficient algorithm for this parametric bootstrap approach is presented in Appendix B.

### Analysis Pipeline

The analysis is performed via an automated bioinformatics pipeline^51^. The inputs are phased variant call format files (VCFs) and a pedigree-based recombination map. Samples are assumed to come from a panmictic population and to be unrelated. The procedure is as follows:

1. Detect IBD segments ≥ 2.0 cM along the genome.
2. Scan for loci where IBD rates exceed four standard deviations above the median.
3. At loci of excess IBD rate,

a. Infer outgroup Â_1_ and the rest of the sample *B*^^^_1_ at the location of the maximum IBD rate.
b. Calculate *Q*_*w*_ for sliding windows of size 250 kb and step 50 kb using Â_1_, *B*^^^_1_. These windows span all positions at which the IBD rate exceeds four standard deviations above the median.
c. Infer outgroup Â_2_ and the rest of the sample *B*^^^_2_ at the location argmax_w_ *Q*_*w*_.

i. Compute IBD entropy given Â_2_ and *B*^^^_2_.
d. Compute *Z*_*j*_ for markers *j* based on Â_2_ and *B*^^^_2_. These scores are calculated for each SNP within 150 kb of the refined location argmax_w_ *Q*_*w*_.
e. Estimate the current frequency of the sweeping allele.
f. Estimate the selection coefficient conditional on the allele frequency estimate.
g. Estimate confidence intervals via parametric bootstrap.

We provide a Python package for the statistical inference of the selection coefficient.

### Demographic Scenarios for Simulation

We consider three demographic models in our simulation studies. We refer to these demographic scenarios as population bottleneck (BN), three phases of exponential growth (G3), and constant size of twenty-five thousand diploids (C25). These parameters refer to the effective population sizes *N*_*e*_(*t*).

The BN model has an ancestral size of ten thousand diploids. Beginning three hundred generations ago, the population experiences two percent exponential growth. At twenty generations ago the effective size instantaneously drops to one million diploids, after which there is continued two percent exponential growth. This bottleneck is a sixty-two percent decrease to the population size. The G3 model has an ancestral effective size of 5,000 diploids. Beginning three hundred, sixty, and ten generations ago, the population grows at exponential rates one, seven, and fifteen percent. These models resemble estimates of *N*_*e*_(*t*) in studies on European American and White British samples^52,53^.

### Simulation of True IBD Tracts from the Coalescent

Using simulated IBD segments greater than 3.0 cM, we assess the empirical properties of our selection coefficient inference. We simulate these segments using the bootstrap algorithm described in Appendix B. Default simulation settings are *s* = 0.030, *p*(0) = 0.50, the BN demographic model, and multiplicative selection. The sweep is incomplete and from *de novo* mutation. The sample size is five thousand diploids.

We change one simulation parameter at a time and explore how the estimator performs.

First, we estimate the selection coefficient conditional on the true allele frequency, effective sizes, and genetic model. We consider the following choices: *s* ∈ [0.02,0.03,0.04]; *p*(0) ∈ [0.25,0.50,0.75]; and the BN, G3, and C25 demographic scenarios. We specify the perturbed model parameter for each row in Table 1. Second, we estimate the selection coefficient conditional on model misspecifications: allele frequency or *N*_*e*_(*t*) different from simulation, and sweeps from variants at one, two, and five percent frequency (abbreviated SV for standing variation). For misspecified demography, 1.1x and 0.9x denote uniform scaling of effective sizes *N*_*e*_(*t*). We specify the true simulated model and the model assumed for estimation for each row in Supplementary Table 2.

**Table 1.** Selection coefficient estimates based on true IBD tracts Summary statistics are the mean estimate, the mean bias, the width of confidence interval, and the percentage of intervals containing the true selection coefficient. Ninety-five percent confidence intervals are based on fifty bootstraps. Each row represents the results of 10,000 replicates. Default settings are *s* = 0.03, *p*(0) = 0.50, a population bottleneck demography, the multiplicative genetic model, an ongoing sweep from a new beneficial mutation. Demographic scenarios considered are population bottleneck (BN), three phases of exponential growth (G3), and constant *N*_*e*_ =25000 (C25). The sample size is five thousand diploids.

| Scenario | True $s$ | Estimate $\bar{\hat{s}}$ | Width | Coverage |
| --- | --- | --- | --- | --- |
| $s = 0.020$ | 0.0200 | 0.0202 | 0.0085 | 97.67% |
| $s = 0.030$ | 0.0300 | 0.0300 | 0.0085 | 95.21% |
| $s = 0.040$ | 0.0400 | 0.0401 | 0.0099 | 95.11% |
| $p(0) = 0.25$ | 0.0300 | 0.0300 | 0.0087 | 95.31% |
| $p(0) = 0.50$ | 0.0300 | 0.0300 | 0.0066 | 94.89% |
| $p(0) = 0.75$ | 0.0300 | 0.0300 | 0.0058 | 95.19% |
| $N_e(t) = \text{BN}$ | 0.0300 | 0.0300 | 0.0066 | 95.26% |
| $N_e(t) = \text{SE}$ | 0.0300 | 0.0300 | 0.0062 | 95.01% |
| $N_e(t) = \text{C25}$ | 0.0300 | 0.0300 | 0.0066 | 95.79% |

We compare the selection coefficient estimate to the truth in terms of bias and the width and coverage of ninety-five percent confidence intervals (from fifty parametric bootstraps). Bias is the estimate subtracted by the true parameter. Width is the (absolute) difference between the left and right endpoints of the confidence interval. Coverage is the proportion of confidence intervals that contain the true *s*. These summary statistics are averaged over 10,000 replicates.

### Simulation of Inferred IBD Tracts from Sequence Data

We investigate the performance of our methods to identify selected alleles, estimate the frequency and location of an unknown sweeping allele, and infer its selection coefficient. We use SLiM^54,55^ and msprime^56^ to generate 8 cM of genetic data for five thousand diploid individuals.

We consider constant mutation and recombination rates of 1e-8, gene conversion rate 2e-8, mean gene conversion tract length of three hundred basepairs, and genotyping error rate 0.02%. At the coalescent time specified in Table 2 and Supplementary Table 3, we place a *de novo* sweep mutation at the center of an 8 cM genomic region. We focus on partial sweeps, so we re-run simulations if the *de novo* mutation is not segregating at a present-day allele frequency between ten and ninety percent. Analyses are based on five thousand diploid individuals sampled from the present-day population. We use true haplotype phase for IBD segment detection. We simulate data one hundred times for each selection coefficient and mutation time. Unless otherwise specified, these are the default settings for simulation.

**Table 2.** Selection coefficient estimates based on IBD segments inferred from sequence data For each row, estimates Ŝ, mean absolute deviation (MAD), and coverage are averaged over one hundred simulations. Ninety-five percent confidence interval is based on fifty bootstraps. Time refers to generations ago when the *de novo* mutation arose. The sample size is five thousand diploids. Population bottleneck (BN) is the demographic scenario. Mutation, recombination, and gene conversion rates are 1e-8, 1e-8, and 2e-8.

| Time | True $s$ | Estimate $\hat{s}$ | MAD | Coverage |
| --- | --- | --- | --- | --- |
| 200 | 0.040 | 0.0407 | 0.0025 | 98 |
| 250 | 0.040 | 0.0383 | 0.0026 | 84 |
| 225 | 0.035 | 0.0361 | 0.0024 | 99 |
| 275 | 0.035 | 0.0336 | 0.0020 | 86 |
| 250 | 0.030 | 0.0304 | 0.0018 | 98 |
| 300 | 0.030 | 0.0295 | 0.0016 | 86 |
| 275 | 0.025 | 0.0264 | 0.0022 | 94 |
| 325 | 0.025 | 0.0254 | 0.0013 | 94 |
| 250 | 0.020 | 0.0220 | 0.0032 | 90 |
| 300 | 0.020 | 0.0216 | 0.0023 | 95 |
| 325 | 0.015 | 0.0168 | 0.0046 | 83 |
| 375 | 0.015 | 0.0175 | 0.0038 | 82 |
| 350 | 0.010 | 0.0136 | 0.0060 | 86 |
| 400 | 0.010 | 0.0145 | 0.0066 | 78 |

Like the simulation study on true IBD segments, we change one parameter at a time and evaluate performance. We explore the BN, G3, and C25 demographic models for *s* = 0.03, and we study selection coefficients *s* ∈ [0.01, 0.015, 0.02, 0.025, 0.03, 0.035, 0.04] for the BN demography model. We use the true effective sizes to estimate the selection coefficient.

Cryptic population structure contributes to LD and can thus confound selective sweep signals. We consider a setting of two equally sized subpopulations splitting 500, 450, and 400 generations ago and randomly mixing at migration rates forty, ten, and one percent. The effective population sizes follow the BN demographic model; the effective population sizes for each subpopulation are half the effective population sizes. We use these effective population sizes to estimate the selection coefficient.

We also examine the impacts of sample size and mutation, recombination, and gene conversion rates. The sample sizes vary between one, two, and five thousand diploids. Mutation and recombination rates vary between 1e-8 and 2e-8. Gene conversion rates vary between zero and 2e-8. For these studies on mutation, recombination, and gene conversion, we simulate sixty replicates. In the case of zero gene conversion, we compare selection coefficient estimates between inferred IBD tracts and true IBD tracts derived from tskibd^57^.

As before, we report for average selection coefficient estimates, the mean absolute deviation between estimate and truth, and the coverage of confidence intervals. Confidence intervals are based on two hundred parametric bootstraps. Estimated coverage is less precise because there are fewer replicates (one hundred instead of ten thousand simulations).

### Comparing Methods for Identifying Selected Alleles

Numerous methods exist to rank plausible sweep variants. We compare iSWEEP against iSAFE^24^, iHS^7^, and nSL^23^. Their inputs are phased genetic data 1.2 Mb to the left and right of the sweeping allele; the input to iSWEEP is genetic data for the entire 8.0 cM region. iHS and nSL are implemented using the selscan software^14^. Default settings are used for all methods. The data input and algorithm settings for each method are summarized in Supplementary Table 4.

We evaluate methods in terms of how well they identify the selected allele or variants strongly correlated with it. The task of identifying the selected allele is far from trivial as there may be tens of thousands of variants in the region of interest. Each method orders the SNPs, where one is the best possible rank estimate. To see if the ordering prefers SNPs in strong LD with the selected allele, we look at mean absolute deviations (MAD) in frequency and location for the ten SNPs of lowest rank. These metrics discern if a method is favoring alleles on the same haplotypes as the selected allele, in the case that the selected allele is not genotyped.

Our method also estimates the frequency of the sweeping allele even if its identity is unknown. This feature is unique to our method. To assess accuracy, we study absolute deviations between our estimates *p*^(0) and the simulated allele frequencies *p*(0).

### Comparing Methods for Estimating Selection Coefficients

Few methods exist to estimate selection coefficients with modern sequences only. We compare the methods CLUES^30^ and ImaGene^39^ against our estimator. CLUES finds a selection coefficient that optimizes a likelihood for the ancestral recombination graph (ARG). ImaGene^39^ trains a convolutional neural network on simulated data, where inputs are genotype matrices and the response variable is the selection coefficient. To apply these methods, we downsample to five hundred of the five thousand simulated diploids due to limitations in computer memory and runtime. The data input and algorithm settings for each method to estimate selection coefficients are summarized in Supplementary Table 4.

CLUES uses importance sampling from a distribution on ARGs. To sample the posterior distribution of ARGs, we run Relate^33^ with a buffer of 100 kb to the left and right of the selected allele, provide true population effective sizes discretized every ten generations, and run five hundred MCMC samples for the branch lengths. We choose these parameters to provide as much data to CLUES as possible while keeping compute times within 1 day for each replicate. We run CLUES using default parameters and the exact allele frequency and position of the sweep mutation. The demographic scenario is coarsened to piecewise constant *N*_*e*_(*t*) changes for the intervals used in Speidel et al. (2019)^33^. This coarsening of *N*_*e*_(*t*) is to reduce compute time to be within one day.

We generate data for ImaGene using the same biological and population bottleneck demographic scenario as in the simulation study. We use the msms software to simulate large amounts of training data because it is faster than SLiM for our examples^58^. The response variables to learn are selection coefficients discretized every 0.005 between 0.00 and 0.05. In total, we simulate two million replicates uniformly proportioned among the different selection coefficients. We split the samples into 80% training and 20% validation datasets. Otherwise, all details are as in the ImaGene paper^39^. Fitting the neural network model takes 165 hours wall clock time using a 24-core computing node. We do not explore other demographic scenarios and varying mutation, recombination, and gene conversion rates because we would have to fit a new ImaGene model each time.

### Trans-Omics for Precision Medicine Data

We focus on selective sweeps in European ancestry samples from the Trans-Omics for Precision Medicine (TOPMed) project. The sweep model we assume has no population structure or familial relatedness. To address these model misspecifications, we subset the 38,387 whole genome sequences in Browning et al. (2021)^59^. We compute principal components (PCs) using the GENESIS and SNPRelate software programs. Based on data visualization, we determine bounds on PCs 1-4 to define European (EUR) and non-European ancestry groups. We approach clustering this way because there are many admixed samples in the TOPMed data. Supplementary Figure 1 shows that samples in the EUR group represent broad European ancestry with slight geographic differentiation.

As validation, we use the 1000 Genomes and Human Genome Diversity Panel (HGDP) datasets to perform supervised global ancestry inference with ADMIXTURE^42^. The CEU (Utah residents with Northern and Western European ancestry), CHB (Han Chinese in Beijing, China), GIH (Gujarati Indians in Houston, TX), and YRI (Yoruba in Ibadan, Nigeria) samples are used from 1000 Genomes and 61 indigenous American samples are used from HGDP^60,61^. For the EUR group, the global ancestry with respect to the CEU reference is minimum 73%, interquartile range (90%, 97%), and mean 93%. Other continental ancestry groups do not have enough samples for the IBD-based analyses explored here.

We remove samples with kinship coefficients ф greater than 0.125 due to close familial relatedness. How we do this is by forming a graph edge if haplotypes *i, j* have ф_*i*,*j*_ ≥ 0.125 and randomly keeping one sample from each connected component. Kinship is estimated using IBDkin with input IBD segments longer than 2.0 cM^62^.

Following these filters, the EUR ancestry group consists of 13,778 samples. This group consists of 4172 White, 9583 Other, and 23 non-White samples from self-report surveys. From the cohort studies, there are 7682 WHI, 2157 MLOF, 1217 VTE, 1108 BioMe, 976 VUAF, 345 FHS, 287 CCAF, and 1 HyperGen samples in the EUR group. Study abbreviations are in the Acknowledgements section.

We conduct a combined analysis and individual analyses for cohorts with sample sizes around one thousand or more. To convert physical distance to genetic distance, we use the 2019 recombination map from deCODE Genetics^63^. Haplotype phasing was applied to all 38,387 samples using Beagle v5.2 without a reference panel as previously described in Browning *et al.* (2021)^59^. In IBD calling, we use the same algorithm parameters as in our simulation study, except we specify the error rate err=1.5e-4 based on a preliminary ibd-ends analysis on chromosomes 19 to 22. To infer the nonparametric effective sizes *N*^^^_*e*_(*t*), we use inferred IBD segments ≥ 2.0 cM as input to IBDNe with default settings^52^. To estimate the selection coefficient, we condition on these effective size estimates *N*^^^_*e*_(*t*) and the allele frequency estimate *p*^(0) from our analysis pipeline. Unless otherwise specified, we assume the additive genetic model 1: (1 + *s*): (1 + 2*s*).

## Results

### Selection Coefficient Estimation Given True IBD Tracts

For this study, we estimate the selection coefficient Ŝ using true IBD tract lengths simulated from the coalescent model under selection. Table 1 shows inference results across a range of scenarios, conditional on correctly specified *p*(0) and *N*_*e*_(*t*). For the BN demographic model and *p*(0) = 0.5, average estimates over ten thousand simulations are roughly equal to the true *s* ∈ [0.02, 0.03, 0.04]. For *s* = 0.03, the estimator is likewise accurate if allele frequency *p*(0) ∈ [0.25, 0.75] or under alternative demographic models. Average widths of ninety-five percent confidence intervals are less than 0.01 for all experiments. Moreover, the ninety-five percent confidence intervals contain the true selection coefficient roughly ninety-five percent of the time. Overall, these results demonstrate that the estimator has empirical behavior expected of a maximum likelihood estimator. The estimator performs robustly in the face of some model misspecifications as well.

Supplementary Table 2 shows inferences results conditional on misspecified *p*(0) or *N*_*e*_(*t*) or a selective sweep from an existing low frequency allele. If the allele frequency used for estimation differs from the true value by 0.10, the average estimates can be slightly biased. This bias affects the coverage of confidence intervals, but here the confidence intervals retain at least ninety percent coverage. Effective sizes misspecified by a scalar factor affect bias and coverage similarly. For example, to fit IBD counts to a misspecified model with elevated effective sizes, a greater selection coefficient is required, and vice versa for lower effective sizes.

Interestingly, the estimator is robust to sweeps from low frequency standing variation. The IBD tracts ≥ 3.0 cM mainly come from common ancestors within the last one hundred generations. As a result, our selection coefficient estimation is learned from recent allele frequency changes. We explore standing variation of one, two, and five percent frequency with selection beginning more than one hundred generations ago. The IBD-based approach cannot distinguish between sweeps from low frequency standing variation versus from *de novo* mutation.

### Identifying the Sweeping Allele in Simulated Sequence Data

iSWEEP and iSAFE show improved ability to pinpoint the causal sweep mutation compared to iHS and nSL (Supplementary Figure 2). For iSWEEP and iSAFE the median rank estimate is nearly optimal, and the ninetieth percentile rank estimate is below thirty among thousands of SNPs. These two methods also assign high scores to SNPs that have frequencies within 10% and physical locations within 250 kb of the adaptive allele.

Figure 1 shows results for the same three metrics but aggregated by either *s* ∈ [0.015, 0.02, 0.025, 0.03, 0.035] or allele frequency bins and *s* = 0.03. Both methods demonstrate nearly optimal median rank estimates for all selection coefficients and allele frequencies below seventy percent. The top ten ranked SNPs from the methods are mostly within ten percent frequency and 250 kb away of the causal mutation.

**Figure 1.**
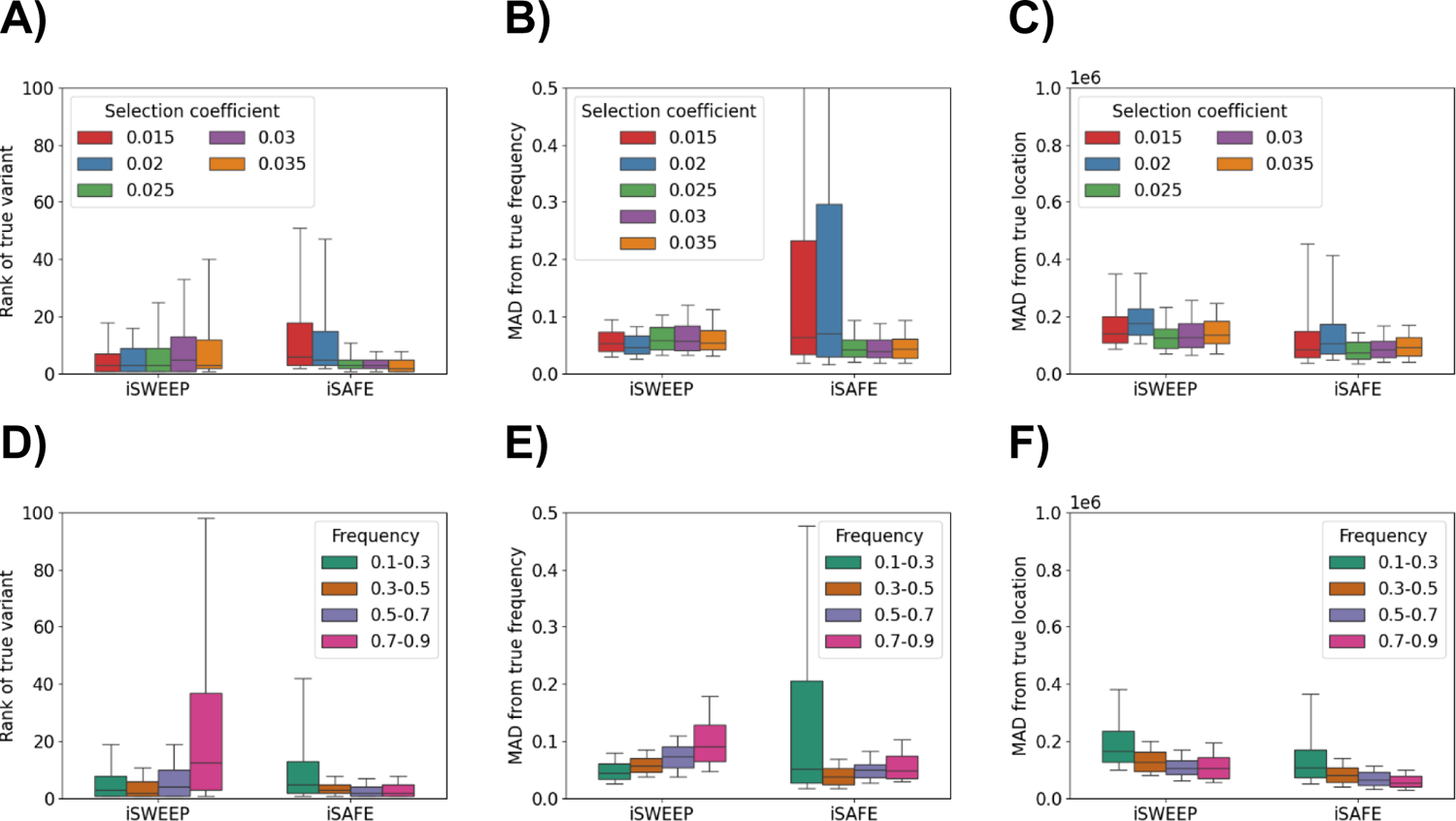
Identifying the sweeping allele and those alleles correlated with it Comparing performance among methods iSWEEP and iSAFE: (A,D) ranking of the true simulated allele, where one is the best possible rank; (B,E) mean absolute deviation (MAD) in frequency over the ten best-ranked alleles; (C,F) MAD in location over the ten best-ranked alleles. (A-C) Results aggregated over different selection coefficients versus (D-E) over different allele frequencies. Box plots show 10^th^, 25^th^, 50^th^, 75^th^, and 90^th^ percentiles for each setting. Two hundred simulations are run for each selection coefficient. The sample size is five thousand diploids. Population bottleneck (BN) is the demographic scenario. Mutation, recombination, and gene conversion rates are 1e-8, 1e-8, and 2e-8.

Both iSWEEP and iSAFE perform well in all simulations, so the following discussion concerns minor differences in accuracy. The ninetieth percentile rank estimates from iSWEEP are less than twenty for *s* ≤ 0.02 but as high as fifty from iSAFE; the ninetieth percentile rank estimates from iSWEEP are as high as forty for *s* ≥ 0.03 but less than fifteen from iSAFE. For *s* ≤ 0.02, iSAFE can sometimes rank SNPs among the top ten that are more than 450 kb away from the causal mutation and off by more than fifty percent from the true frequency.

When the sweeping allele is at frequencies greater than seventy percent, the median and ninetieth percentile rank estimates from iSWEEP are nearly as high as twenty and one hundred.

When the sweeping allele is at frequencies lower than thirty percent, iSAFE may rank SNPs among the top ten that segregate at frequencies more than fifty percent removed from the true frequency. iSWEEP outperforms iSAFE when the sweeping allele frequency is less than seventy percent. This result is not surprising because iSWEEP is based on IBD derived from a very recent adaptive haplotype.

Supplementary Figure 3 suggests that both methods are robust to the BN, G3, and C25 demographic scenarios. iSWEEP manages the BN demographic scenario better whereas iSAFE manages the G3 scenario better. The performance of the IBD outgroup algorithm used by iSWEEP is driven by its main data input: abundant and accurately inferred IBD segments. For BN and C25 demographic scenarios, there are more IBD tracts to learn from compared to the G3 demographic scenario.

We also explore the performance of iSWEEP in cases of smaller sample size, different mutation, recombination, and gene conversion rates, and cryptic population substructure.

Supplementary Figure 4 demonstrates that iSWEEP’s performance improves with increasing sample size. Performance may be especially poor for sample sizes less than one thousand diploids, in which case there is often less than one hundred IBD segments in the BN demographic scenario simulations. Supplementary Figure 5 shows similar summary statistics for zero versus 2e-8 gene conversion rates. This result may indicate that the ibd-ends tool properly addresses small gene conversion tracts in sequence data. Likewise, we find indistinguishable results in Supplementary Figure 6 in different combinations of mutation and recombination rates. Finally, Supplementary Figure 7 shows the potential of population substructure to impact accuracy in identifying the sweeping allele. Rank estimates for the sweeping allele get much worse with stronger population structure or if the population split is concurrent with the beginning of the sweep.

Variant scores in iSWEEP have clear interpretations and require few assumptions. The scores are simple differences in proportions. They do not require prior knowledge of which allele is favored, but instead infer the favored allele as the one at higher frequency in the outgroup of excess IBD.

The IBD entropy statistic measures cluster agglomeration, where zero indicates one large outgroup. We observe that IBD entropy is greater than 0.50 for more than twenty percent of BN demographic scenario simulations with allele frequency less than thirty percent. On the other hand, IBD entropy is zero for all BN demographic scenario simulations with allele frequency greater than thirty percent. This finding indicates that IBD entropy less than 0.50 may be sufficient evidence to support the hard sweep model if the sweeping allele frequency is intermediate to high. IBD entropy less than 0.50 is not necessary evidence for sweeps presently at low frequency, in which case the rank estimates can be very accurate (Figure 1). An explanation for IBD entropy greater than 0.50 could be that the locus has multiple adaptive alleles, violating the sweep model.

To model a selective sweep, we require an allele frequency estimate, not the identity of the sweeping allele. Only our method provides such an allele frequency estimate. We aggregate information over best-ranked SNPs to estimate *p*^(0). Supplementary Figure 8 compares the absolute deviation between this estimate and the true frequency. For *p*(0) ≤ 0.7, the median and ninetieth percentiles for the absolute deviation between *p*(0) and *p*^(0) are roughly within 0.05 and 0.10. Supplementary Table 2 shows that selection coefficient estimation can be slightly biased if the allele frequency estimate is off by 0.10. Specifically, for *s* = 0.03, we observe average bias in Ŝ less than 0.0005 when *p*(0) = 0.50 and *p*^(0) = *p*(0) ± 0.10 and less than 0.002 when *p*(0) = 0.25 and *p*^(0) = *p*(0) ± 0.05.

The accuracy of our estimates *p*^(0) are robust to different selection coefficients and the three demographic models considered (Supplementary Figure 8). The estimates are robust to different mutation, recombination, and gene conversion rates as well (Supplementary Figure 5 and 6). Estimating *p*(0) can be unsatisfactory in examples of *p*(0) ≥ 0.7 or cryptic population substructure (Supplementary Figure 7). Estimating *p*(0) can also be inaccurate with the small sample size of one thousand diploids (Supplementary Figure 4).

### Selection Coefficient Estimation Given Inferred IBD Tracts

Given an allele frequency estimate and accurate IBD detection, we expect inference of Ŝ to be in line with those of the idealized coalescent simulation study. Table 2 and Supplementary Table 3 present average bias in selection coefficient estimation and confidence interval coverages even when the allele frequency is estimated and inferred IBD tracts are used. We observe average bias less than 0.002 for *s* > 0.02 cases; we observe more bias in our estimator for *s* ≤ 0.02. This bias is not due to inaccurate IBD detection (Supplementary Figure 9), but rather that IBD rates are similar between neutral evolution and weak selection. These weak sweeps may not pass our genome-wide selection scan. Generally speaking, confidence interval coverage is greater for the more recent times of *de novo* mutation and larger selection coefficients. Compared to the results in Table 1, the decrease in coverage can be attributed to conditioning on biased *p*^(0).

We contrast our estimation procedure with the deep learning method ImaGene^39^ and the tree inference method CLUES^30^. Figure 2 reports results of this comparison for varying selection coefficients and demographic models. The tenth and ninetieth percentiles of our estimates contain the true selection coefficient for *s* ≥ 0.015; the interquartile range of our estimates contain the true selection coefficient for *s* ≥ 0.025. These findings apply to the C25 and G3 models as well when *s* = 0.03. Estimates of *s* from ImaGene are consistently inflated and estimates of *s* from CLUES do not increase when we increase the true selection coefficient. In fact, estimates from CLUES are around or below 0.01 for most simulations. The accuracy of our estimator carries over to examples of smaller sample size, cryptic population substructure, and varying mutation, recombination, and gene conversion rates as well (Supplementary Figures 4, 5, 6, and 7).

**Figure 2.**
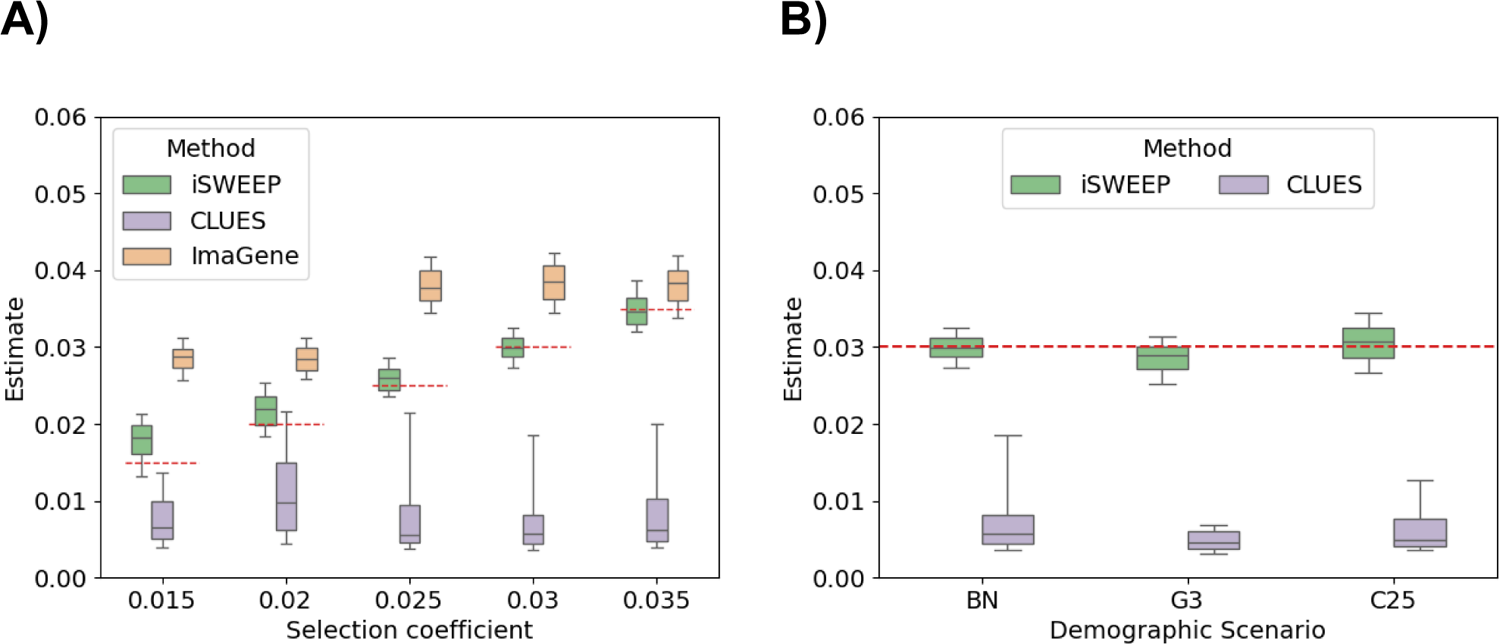
Estimating selection coefficients from simulated sequence data Comparing performance among methods iSWEEP, CLUES, and ImaGene: (A) fixed population bottleneck (BN) effective sizes, varying selection coefficients; (B) fixed selection coefficient *s* = 0.03, varying demographic scenarios: population bottleneck (BN), three phases of exponential growth (G3), and constant size *N*_*e*_ = 25,000 (C25). Box plots show 10^th^, 25^th^, 50^th^, 75^th^, and 90^th^ percentiles for each setting. Two hundred simulations are run for each selection coefficient. Horizontal dashed red lines correspond to the true selection coefficient. The sample size is five thousand diploids for iSWEEP. Due to memory and runtime limitations, the sample size is five hundred diploids for CLUES and ImaGene. Mutation, recombination, and gene conversion rates are 1e-8, 1e-8, and 2e-8.

The authors of CLUES acknowledge that their method is limited to *s* ≤ 0.01, and they only test performance on simulations where *s* ≤ 0.01 ^30,31^. An analysis of CLUES from separate authors observes results like ours for 0.01 ≤ *s* ≤ 0.02. On the other hand, they report that CLUES can be accurate in this range when given the true simulated tree^38^. Knowing the true coalescent tree is unrealistic for any analysis. One explanation for the results we observe for CLUES is that inferring the high-dimensional ARG is extremely difficult^64^. ARG inference may depend on the ratio between mutation and recombination rates as well^31^, while iSWEEP only requires accurate IBD calling. Vaughn and Nielsen (2023) show that using ancient DNA in the true ARG can improve the CLUES inference^31^. Overall, we suggest our method as suited for recent strong positive selection *s* ≥ 0.015 whereas current alternatives address weak selection *s* ≤ 0.015 ^30,31,38,39^.

### Selection Scan in TOPMed Samples of European Ancestry

Table 3 summarizes the results of our selection scan in the TOPMed cohorts. In the combined EUR group, we find eight regions where IBD rates exceed four standard deviations above the genome-wide median and three or more cohort-specific analyses replicate the signal. The inconsistencies between analyses apply to the selection signals which marginally pass our genome-wide threshold. Seven of these eight consensus loci appear in prior selection scans on White British individuals in the UK Biobank^8–10^ and pan-European analyses using ancient DNA^15,26^. The signal between 205 to 207 Mb on chromosome 1 appears in the combined EUR, WHI, and MLOF groups, but it does not appear in the Browning and Browning (2020) study on a White British cohort^8^; the IBD rates in this region barely exceed four standard deviations above the genome-wide median.

**Table 3.** Eight regions highlighted in selection scan on TOPMed European Americans Loci where identity-by-descent (IBD) rate exceeds four standard deviations above the genome-wide median, and at least three out of six cohort studies pass all criteria given in the main text. Physical and genetic positions for the location of maximum IBD rate are shown in megabases (Mb) and centiMorgans (cM). The intervals denote the range over which IBD rates exceed the scan threshold. Locations are aligned to build hg38 and the 2019 pedigree-based recombination map from deCODE genetics. Genes of interest are annotated for regions discussed in the main text. Rows are based on analysis of the 13,778 samples in the combined EUR group, except the row for OCA2 is based on the 7,682 samples in the WHI cohort.

| Chr | Max IBD Rate<br>(1e-4 scale) | cM | Position (Mb) | # studies | Genes |
| --- | --- | --- | --- | --- | --- |
| 2 | 9.09 | 141.79 | 134.84 (132.52-139.90) | 6 / 6 | LCT |
| 6 | 7.20 | 49.35 | 30.80 (24.12-36.12) | 6 / 6 | MHC |
| 15 | 3.31 | 15.42 | 31.18 (30.34-32.16) | 6 / 6 | TRPM1 |
| 12 | 2.72 | 120.54 | 113.08 (110.90-113.64) | 5 / 6 | OAS1-2-3 |
| 6 | 2.14 | 107.11 | 105.98 (103.76-108.46) | 3 / 6 | PRDM1 |
| 1 | 2.13 | 205.05 | 206.62 (205.50-207) | 3 / 6 | . |
| 5 | 2.06 | 53.34 | 33.96 (31.00-35.98) | 3 / 6 | SLC45A2 |
| 15 | 2.22 | 10.19 | 28.10 (25.94-30.86) | 3 / 6 | OCA2 |

Figure 3 shows the IBD rates along the autosomes, the median, and the threshold for the EUR group. Supplementary Figure 10 exhibits similar results among the five cohorts. The four largest signals at LCT, MHC, TRPM1, and OAS are thirty-five, twenty-six, ten, and eight standard deviations above the population median IBD rate, respectively. All six analyses observe excess IBD at LCT, MHC, and TRPM1, and five of them observe excess IBD at OAS.

**Figure 3.**
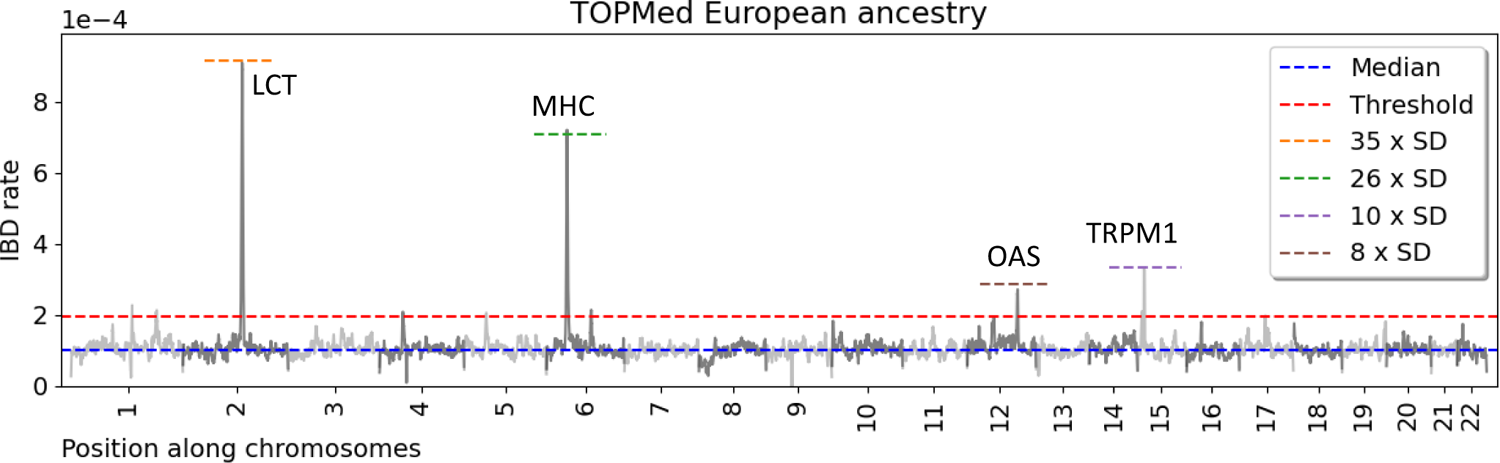
Selection scan in TOPMed Americans of European ancestry (EUR) Identity-by-descent (IBD) segments longer than 2.0 cM were inferred using hap-ibd^17^ and ibd-ends^8^. The rate of IBD in the population sample is computed every 20 kb. The median (blue) plus four times the standard error (red) are presented as horizontal lines. LCT, MHC, OAS, and TRPM1 loci are annotated, and horizontal lines give the number of standard deviations above the population median.

### Modeling LCT Selection

Lactase persistence is widely believed to be subject to a selective sweep in European ancestry populations. We view it as a positive control to evaluate our method. The putative selected allele is a regulatory mutation −13.910C>T upstream of LCT (OMIM: 603202) in intron 13 of MCM6 (OMIM: 601806). This allele enhances the promoter of LCT, allowing an alternative path for gene expression.

Here the individual and aggregate analyses show nearly perfect concordance. IBD entropies are zero, and there are single large IBD outgroups that comprise fifty percent or more of the samples. Five of six analyses rank the −13.910C>T mutation as the maximally differentiated SNP between the inferred outgroup and the rest of the sample. Supplementary Figure 11A depicts scores for the SNPs flanking the causal mutation. Our allele frequency estimate without prior knowledge of the selected allele is *p*^(0) = 0.725, while the frequency of −13.910C>T is 0.6864. We estimate the LCT selection coefficient using the frequency of the consensus −13.910C>T mutation.

The lactase persistence phenotype is dominant but the enzymatic activity is additive^26^. What phenotype is selected for and which form of genic selection is never known. Given the additive model, we estimate Ŝ_a_ = 0.0325 (95% *CI* = (0.0278, 0.0373)). Supplementary Table 5 gives the additive model estimates for BioMe, MLOF, VTE, VUAF, and WHI cohorts. Given the dominance model, we estimate Ŝ_d_ = 0.0488 (95% *CI* = (0.0423, 0.0552)).

Figure 4A shows the historical allele frequencies implied by Ŝ_a_and Ŝ_d_. To display historical allele frequencies, we draw parametric bootstraps over selection coefficients Ŝ_b_ and simulate allele frequencies backwards in time with selection and genetic drift. This boostrap represents uncertainty in the IBD process and genetic drift. Mathieson and Terhorst (2022)^26^ and Vaughn and Nielsen (2023)^31^ model LCT selection with time-varying coefficients. Their approaches use ancient DNA; our method does not leverage temporal information, and thus estimating time-varying selection coefficients is out of its scope. However, the historical allele frequencies from our dominance model more closely match the curve in Mathieson and Terhorst (2022)^26^ than the historical allele frequencies from our additive model. Compared to the other loci, the sweep at LCT may have progressed more rapidly than any other putative sweeps in Europeans.

**Figure 4.**
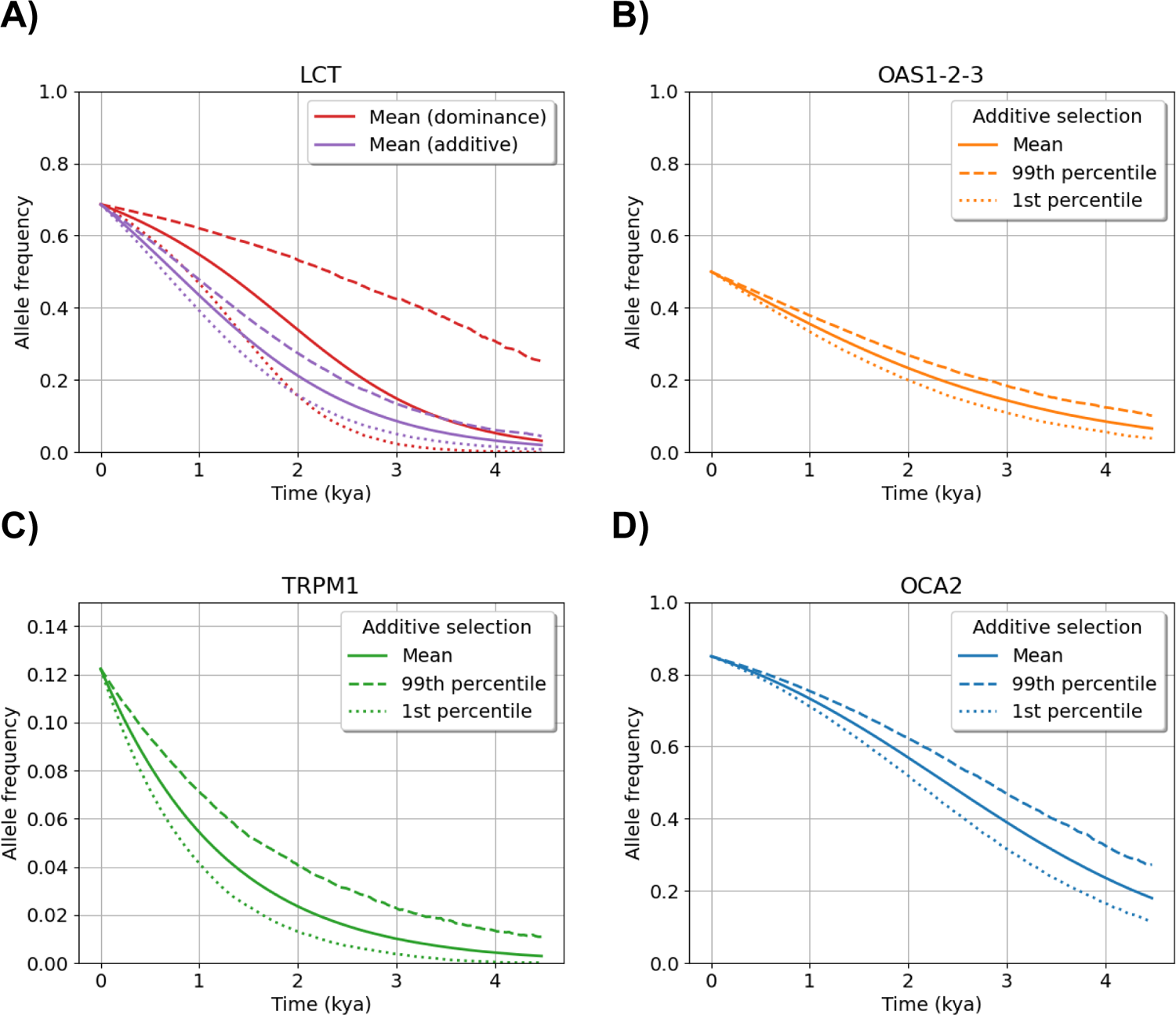
Estimated historical allele frequencies in last 150 generations for four human loci under selection The plots show the means, the first percentiles, and the ninety-ninth percentiles for allele frequencies *p*(*t*) from one thousand parametric bootstraps over identity-by-descent and Wright-Fisher processes for selective sweeps at (A) LCT, (B) OAS 1-2-3, (C) TRPM1, and (D) OCA2 loci. LCT and TRPM1 are based on the best ranked SNP because there is consensus across cohort analyses. OAS1-2-3 and OCA2 use haplotype-based frequency estimates because there is no consensus on a variant. LCT, OAS1-2-3, and TRPM1 are based on IBD segments in the combined EUR group; OCA2 is based on IBD segments in the WHI cohort, the largest cohort where this locus passes the selection scan. Additive selection is assumed, but dominance selection is also shown for LCT. The *y*-axis scale for the (C) TRPM1 locus is changed for improved visibility. Time is measured in thousands of years ago (kya) using one generation equals thirty years.

### Possible Selective Sweeps on Introgressed Haplotypes in OAS 1-2-3 Genes

Studies on archaic introgression have identified a Neanderthal haplotype spanning the OAS genes 1, 2, and 3 (OMIM: 164350, 603350, 603351). These genes transcribe antiviral proteins and may be an example of selection for immune response^65,66^. The introgressed haplotype harbors a splice variant that upregulates expression of a specific isoform^67^. In our six analyses, the locations of maximum IBD rate are 1.5 Mb to the left of OAS1-2-3, instead lying within the LINC02356 (Ensembl: ENSG00000257595), SH2B3 (OMIM: 605093), and ATXN2 (OMIM: 601517) genes. Supplementary Figure 11B shows a long haplotype stretching between these genes all the way to the OAS genes around chr12:113,000,000.

Applying our estimation methods, we arrive at present-day frequency estimates between 0.40 and 0.50. Prior studies have estimated the introgressed Neanderthal haplotype to have a frequency of approximately forty percent among modern Europeans^65–67^. In Supplementary Table 5, three of the cohort analyses rank as first the SNP at basepair chr12:111,270,654. This marker is in the CUX2 gene (OMIM: 610648). IBD entropies are roughly 0.50 across our five analyses; these IBD entropies do not rule out OAS1-2-3 as a sweep locus, but there is weaker evidence for a selective sweep than at LCT. For the combined EUR group, we estimate Ŝ conditional on the haplotype-based frequency estimate *p*^(0) = 0.50, which is close to the frequency of consensus best-ranked SNP from three of five studies. Our estimate of additive selection is Ŝ = 0.0182 (95% CI = (0.0156, 0.208)). According to our simulation study, this low selection coefficient estimate could be inflated.

The evolutionarily conserved haplotypes in the OAS1-2-3 region differ from our modeling assumptions of a single causal allele. We emphasize that the causal allele in our model is not necessarily a SNP variant but could be a haplotype with high LD. Moreover, introgressed haplotypes come from standing variation, not a *de novo* mutation. Our statistical method can estimate selection coefficients in such a soft sweep (Supplementary Table 2), but the interpretation of adaptive introgression is out of its scope. The implied historical allele frequencies are in Figure 4B. They assume the sweep from *de novo* mutation, but the curve in more recent time periods should not depend on this assumption.

### Possible Selective Sweeps around Pigmentation Genes OCA2 and TRPM1

Alleles in pigmentation genes could improve vitamin D levels at northern latitudes. The chromosome 15 region containing TRPM1 (OMIM: 603576) and OCA2 (OMIM: 203200) appears in our IBD-based selection scan as in a related report^8^. In particular, OCA2 has been studied in numerous selection studies^26,30,38^. These genes express in retinal cells and melanocytes and have been linked to eye and skin color phenotypes^68,69^.

For TRPM1, our analysis in each cohort ranks first the SNP at basepair 31,437,832 (Supplementary Table 5). This is the only example in our work where each of the cohorts ranks best the same SNP. This marker is in a splice variant of the KLF13 (OMIM: 605328). Its frequency ranges from 11-13%, and its location is nearly 300 kb to the right of TRPM1. We perform estimation conditional on the allele frequency of this consensus best-ranked SNP and additive selection. We estimate selection coefficients between 0.026 and 0.030 for all of the five cohort-specific analyses. In the combined EUR group, we estimate Ŝ = 0.0273 (95% CI = (0.0200,0.0350)). The implied historical allele frequencies are in Figure 4C.

The neighboring gene OCA2 passes our selection scans in only three cohorts. This locus does not appear in the combined EUR group analysis because its excess IBD rates span only 0.95 cM. SNPs in this region are known to inhibit gene regulation and affect the blue eye color phenotype^70,71^. In Supplementary Table 5, we compute IBD entropies of zero or below 0.50 for these analyses. However, different SNPs are ranked best for each cohort. Supplementary Figure 11C shows that this region contains ∼200 kb of low marker coverage upstream of the rs12913832 variant discussed in Visser et al. (2012)^71^. Low marker density as well as small sample size can impact the precision in our methods, instead emphasizing hitchhiking variants further from the selected allele. On the other hand, these intermediate frequency SNPs at basepairs chr15:28,100,878 and chr15:28,141,480 may be functionally important to an unknown selected haplotype. Since there is not a consensus best-ranked variant, we use the haplotype-based frequency estimate *p*^(0) = 0.85 from the WHI cohort. (The frequency of the rs12913832 variant is 78% in the WHI sample.) We estimate the additive selection coefficient Ŝ = 0.0225 (95% CI = (0.0196,0.0254)). The implied historical allele frequencies are in Figure 4D.

### Limited Evidence for the Sweep Model at Many Human Loci

MHC contains human leukocyte antigen genes that encode molecules which bind to antigens and mediate the presentation of these to the surface of immune cells. These proteins help cells to recognize self from foreign, thereby influencing donor compatibility for organ transplant. IBD entropies at MHC are below 0.50 for the small cohorts but above 0.50 for WHI and the combined EUR group. Additionally, we infer locations for the sweep that differ by as much as seven Mb across studies. These IBD entropies and location estimates indicate that there may be multiple sweeps or multiple adaptive haplotypes in the MHC region. Balancing selection is a clear violation of our model, as are multiple independent sweeps.

Estimating a selection coefficient with our method involves assuming the sweep model. We do not estimate selection coefficients for other loci in Table 3 because they do not have as clear a signal in our analyses. These loci have IBD entropies greater than 0.50, no consensus best-ranked SNP, and the best-ranked SNPs have vastly different allele frequencies and locations. This list includes the genes PRDM1 (OMIM: 603423) and SLC45A2 (OMIM: 606202), which have been discussed before in other selection scans^8,15,26^. PRDM1 has been identified in genome-wide association studies on autoimmune diseases^72^ and SLC45A2 is involved in skin pigmentation^73^. For SLC45A2, Supplementary Figure 11D illustrates an unusual spectrum of common variation between forty and sixty percent around chr5:34,350,000. This region is 400 kb to the right of SLC45A2 and its putative target SNP rs16891972 ^26^. These loci could be examples of older sweeps, sweeps with small selection coefficients, balancing selection, or other evolutionary processes.

## Discussion

We develop a suite of new methods to model very recent selective sweeps from sequence data in the present generation. Our approach uses inferred identity-by-descent tracts which can be informative about a positively selected locus. Our methods pinpoint selection signals to a resolution of 250 kb, estimate frequencies for unknown sweeping alleles, and infer selection coefficients. The selection coefficient estimator we propose is easy to interpret: more IBD tracts results in a larger estimate.

In a coalescent simulation study, we demonstrate that our selection coefficient estimator is accurate and robust to model misspecifications. In simulation studies on sequence data, we compare our methods with existing methods. We observe our approach to identify the sweeping allele to be competitive with state-of-the-art methods. We also find our selection coefficient estimation to be favorable to coalescent and machine learning-based methods for strong selection coefficients *s* ≥ 0.015.

As a case study, we apply our methods to study positive selection in TOPMed European Americans. We estimate the selection coefficient for signals at the LCT, OAS1-2-3, TRPM1, and OCA2 genes. We also identify variants near the TRPM1 and the OAS1-2-3 loci that are strongly differentiated between an inferred IBD outgroup and the rest of the sample. There is limited evidence in our data to fit hard selective sweep models for the MHC, PRDM1, and SLC45A2 loci. Further studies could explore to what extent balancing selection and population structure confound statistical inference of selective sweeps.

For selection at LCT, we estimate ninety-five percent confidence intervals for the selection coefficient that are of width less than 0.01. In comparison, Bersaglieri et al. (2004)^74^ report a confidence interval that has width greater than 0.10. There is a distinction between parameter uncertainty and model uncertainty. The dropout technique in Hejase et al. (2022)^38^ expresses model uncertainty, meaning that we cannot interpret their confidence intervals in a strict probabilistic formulation. Their confidence intervals of width less than 0.001 may be too small to cover the true selection coefficient. Likewise, the high density posterior intervals in Torada et al. (2019)^39^ express uncertainty in the ImaGene model by Monte Carlo sampling from the trained distribution. The Bayesian approach of Stern, Wilton, and Nielsen (2019)^30^ could offer parameter uncertainty by sampling from the posterior distribution, but to get a single estimate their method runs for more than one day in our simulation study.

An estimator should express uncertainty, for instance via a confidence interval. The existing coalescent and machine learning-based methods discussed here do not provide confidence intervals. Our ninety-five percent confidence intervals contain the true selection coefficient in ninety-five percent of our simulations. Compared to prior methodological work, we study cases of strong selection where *s* ≥ 0.01. While our selection coefficient estimation is biased for *s* ≤ 0.015, an IBD-based selection scan can filter loci to just putatively strong sweeps^8^. The confidence interval coverages we observe in simulation study are adequate for *s* ≥ 0.02 even when the sweeping allele is unknown and IBD tracts are inferred.

Our methods are not a panacea for all selection studies in all populations and species. One requirement of our inference is accurate detection of IBD tracts. Pedigree-based genetic maps are preferred, as the selection scan could result in many false discoveries with LD-based maps^8^. Additionally, we use these IBD tracts to infer effective population sizes, which are conditioned on to estimate the selection coefficient. Estimates of recent effective population sizes can be precise for some human populations^52,53,75^. We do not know if the threshold in our selection scan controls the false discovery rate when adjusting for multiple testing. IBD rates across sliding windows can contain the same IBD segments, making the derivation of a genome-wide significance threshold complicated.

All the results are obtained from automated bioinformatics pipelines that run the statistical methods presented here. These methods are implemented in the freely available iSWEEP software package. Algorithm settings can be specified in a configuration file to reproduce the results in this work. The most computationally intensive aspects of our methods involve the use of efficient hap-ibd and ibd-ends algorithms to infer IBD tracts^8,17^. Using a 24-core Intel Xeon Silver 4214 2.2 GHz compute node, wall clock computing time to detect ≥1.0 cM IBD tracts on chromosome 2 for 13,778 samples is 4.5 hours for hap-ibd and 12.5 hours for ibd-ends. These algorithms scale to the analysis of large sequence datasets which far exceed the sample size limitations of existing approaches. The tool ibd-ends is especially important in order to adjust for genotyping errors, gene conversion, and regions of low marker density in sequence data^8^.

Our methods do not need to know which phenotype is being selected for, nor do they require that the sweeping allele be genotyped. The input data is high coverage whole genome sequences from the current generation as opposed to other methods based on ancient or archaic DNA. The IBD-based analyses we conduct on cohorts of less than 2,000 samples are broadly consistent with our analysis on a larger consortium dataset of 13,778 samples. These model assumptions and data requirements may facilitate selection studies on non-European or non-human populations.

## Appendix A: The Identity-By-Descent Model with Selection

Let *p*_*s*_(*t*) and *N*_*e*_(*t*) be the allele frequency of the sweeping allele and the haploid effective size at coalescent time *t* > 0 (generations ago). These functions define subpopulations *N*_*e*_(*t*) × *p*_*s*_(*t*) and *N*_*e*_(*t*) × (1 − *p*_*s*_(*t*)) that have or do not have the sweeping allele. For selection in diploids, we consider four genetic models: additive, multiplicative, dominant, and recessive. These models provide relative fitness between genotypes:

- Additive: (1 + 2*s*): (1 + *s*): 1
- Multiplicative: (1 + *s*)^2^: (1 + *s*): 1
- Dominant: (1 + *s*): (1 + *s*): 1
- Recessive: (1 + *s*): 1: 1

The recursive formulas for these genetic models are as follows:

- Additive: *p*_*s*_(*t* − 1) = *p*_*s*_(*t*) × (1 + *s* + *s* × *p*_*s*_(*t*))/(1 + 2 × *s* × *p*_*s*_(*t*))
- Multiplicative: *p*_*s*_(*t* − 1) = *p*_*s*_(*t*) × (1 + *s*)/(1 + *s* × *p*_*s*_(*t*))
- Dominant: *p*_*s*(*t*−1)_ = *p*_*s*_(*t*) × (1 + *s*)/(1 + 2*s* × *p*_*s*_(*t*) × (1 − *p*_*s*_(*t*)) + *s* × *p*_*s*_(*t*)^2^)
- Recessive: *p*_*s*_(*t* − 1) = *p*_*s*_(*t*) × (1 + *s* × *p*_*s*_(*t*))/(1 + *s* × *p*_*s*_(*t*))

For *p*_*s*_(*t*) in terms of *p*_*s*_(*t* − 1) and *s*, we invert *p*^−1^(*t* − 1) with the quadratic formula and choose the solution in the interval [0,1]. Unless otherwise specified, we assume the additive model. Selection coefficients from different genetic models are not comparable.

These formulas concern the mean behavior in the Wright Fisher process with drift and selection. Let the random count variable *X*(*t*) ∼ *Binomial*(2*N*_*e*_(*t*), *p*_*s*_(*t* − 1)) be the number of haplotypes with the sweeping allele. For estimating *s*, we use the deterministic *p*_*s*_(*t*) from the backward formulas. For bootstrap uncertainty, we use estimates *p*^_*s*_(*t*) = *X*(*t*)/[2*N*_*e*_(*t*)] when simulating back in time. Note that the sample mean estimate *p*^_*s*_(*t*) is a maximum likelihood estimate and should be close to *p*_*s*_(*t*) for large *N*_*e*_(*t*).

Let coalescent time *T* and identity-by-descent (IBD) tract length *L* be random variables. Let *c* be a user-defined genetic distance threshold. Indices *i, j* refer to haplotypes that share IBD tracts from a common ancestor. For two randomly sampled haplotypes,

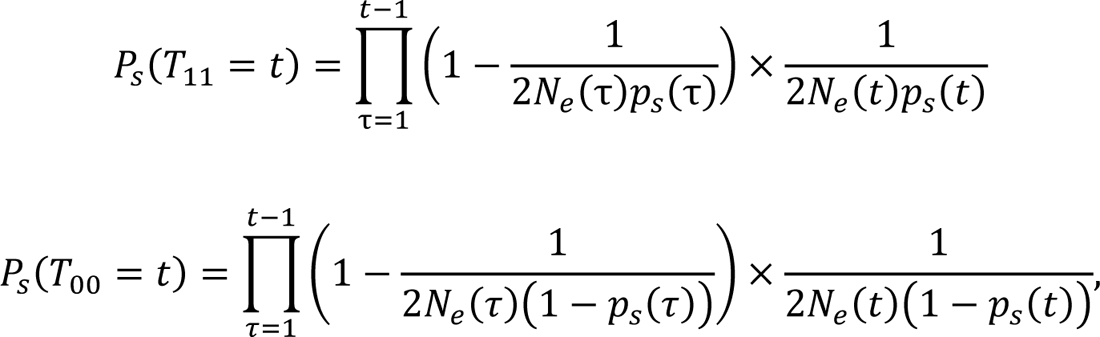

where *T*_11_, *T*_00_ are the coalescent times for randomly sampled haplotypes with alleles 1 and 0. For the biallelic marker, the probability mass for the coalescent time is

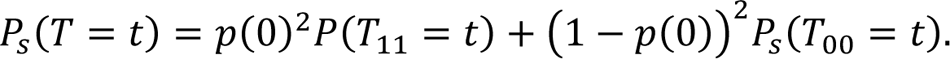

These formulas generalize to the multiallelic setting.

Recombination endpoints to the right of a focal position are exponentially distributed with rate equal to *g*(*t*). We use the function *g*(*t*) = *t*/100 to measure genetic distance in centiMorgans (cM) instead of Morgans. The shared IBD to the right of the focal position from two independent, identically distributed recombination endpoints is the minimum length. Thus, the entire IBD tract length *L* | *T* = *t* ∼ *Gamma*(2, 2 × *g*(*t*)) is the sum of shared IBD to the left and right of the focal position^52,76^. The probability *P*_*s*_(*L* ≥ *c*) = ∫ ∫ *P*_*s*_(*T* = *t*)*P*(*L* ≥ *c*|*T* = *t*) is calculated using the law of total probability.

## Appendix B: Estimation of Selection Coefficients

We discuss three statistical properties of maximum likelihood estimates (MLEs) as they apply to our work. First, consistency means that an estimator Ŝ for the parameter *s* gets arbitrarily close to *s* as sample size *M* increases. Second, sufficiency means that Ŝ uses all relevant information to estimate *s*. Third, MLEs are normally distributed in large samples. Let ^*P*^^_*s*_(*L* ≥ *c*) be an MLE for *P*_*s*_(*L* ≥ *c*), and let *f* be the function that maps *P*_*s*_(*L* ≥ *c*) to *s*. Theorem 7.2.10 from Casella and Berger (1993) says that *f*(^*P*^^_*s*_(*L* ≥ *c*)) is the MLE for the selection coefficient *s*^77^.

Let *L*_*i*,*j*_ be the IBD tract length shared by haplotypes *i* and *j* in a sample of *n*. The binary random variable *Y*_*i*,*j*_ = *I*(*L*_*i*,*j*_ ≥ *c*) denotes if *L*_*i*,*j*_ is longer than threshold *c*. The *Y*_*i*,*j*_’s are identically distributed Bernoulli random variables with probability parameter *P*_*s*_(*L* ≥ *c*), but they are correlated via the ancestral tree. The empirical tail probability 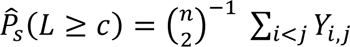 is always an unbiased estimator of *P*_*s*_(*L* ≥ *c*). We want even stronger statistical properties for estimation. Consider the composite likelihood assumption in which (*Y*_1_, *Y*_2_, …, *Y*_*M*_) are *independent*, identically distributed Bernoulli random variables with probability parameter *P*_*s*_(*L* ≥ *c*). The sample mean ^*P*^^_*s*_(*L* ≥ *c*) = Ŷ_1:*M*_ is then the MLE for the binomial experiment. In other words, the IBD rate maximizes the composite likelihood, which typically gives consistent point estimates in statistical genetics^78^.

We propose the value Ŝ that solves the equation 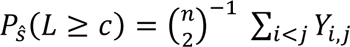 as our estimator. Because inverting *P*_*s*_(*L* ≥ *c*) is analytically intractable, we optimize the expression 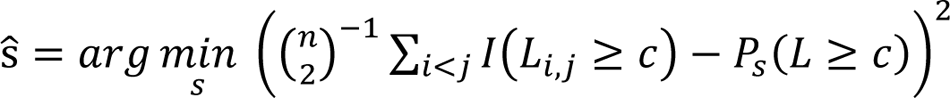. This function *P*_*s*_(*L* ≥ *g*) is not necessarily monotone in *s*, and proving such is intractable for general form *N*_*e*_(*t*) and integration over the recombination endpoints. However, it is related to the probability of IBD alleles, which Albrechtsen, Moltke, and Nielsen (2010) prove to be increasing when selection acts on an allele^11^. Numerically, one can verify monotonicity for *P*_*s*_(*L* ≥ *c*) in *s*, which is the case in all experiments pursued here. Theorem 7.2.10 from Casella and Berger (1993) does not require monotonicity^77^, but the notion of a one-to-one relationship between the IBD rate and the selection coefficient is appealing.

In simulation study, we find that the average Ŝ̅ is close to the true value. In Supplementary Figure 12, we illustrate the inferred counts of IBD tracts ≥ 2.0 cM in the combined EUR group and the empirical distribution of Ŝ given *p*(0) = 0.50 and the estimated effective sizes. Their distributions appear to be normally distributed. These two arguments suggest that our estimator behaves like an MLE. Deep learning methods say that, instead of low-dimensional statistics, we require high-dimensional data to infer selection coefficients ^38,39,79^.

This assertion is not true for sufficient statistics. Under the composite likelihood, the IBD rate is a one-dimensional sufficient statistic for *P*_*s*_(*L* ≥ *c*).

We provide a parametric bootstrap approach to give ninety-five percent confidence intervals^80,81^. The parametric bootstrap draws realizations from a probability model conditional on a parameter estimate, and it then re-estimates the parameter with the bootstrap sample.

Simulating tracts independently undersells variance in the IBD tract length distribution, affecting the width of confidence intervals (Supplementary Figure 13). We present an algorithm to generate IBD segments from the coalescent around a focal site given parameter estimates for the hard sweep.

Algorithm 1.

1. Initialize all nodes *i* to have recombination endpoints *l*_*i*_, *r*_*i*_ = ∞.
2. Simulate a conditional coalescent tree given *s, p*(0), *N*_*e*_(*t*), and a genetic model.
3. At each coalescence to a common ancestor:

a. Draw left and right recombination endpoints *l*_*I*_, *r*_*I*_ ∼ *Exp*((*t*_*k*_ − *t*_*I*_)/100) where *t*_*I*_, *t*_*K*_ is the time ago for some incoming interior node and the common ancestor.
b. Update *t*_*I*_ = *t*_*K*_, *r*_*i*_ = min (*r*_*i*_, *r*_*I*_), *l*_*i*_ = min (*l*_*i*_, *l*_*I*_).
c. Compute *r* = min(*r*_*i*_, *r*_*j*_) and *l* = min (*l*_*i*_, *l*_*j*_) for sample nodes *i, j* under interior nodes *I, J*.
d. If *r* + *l* ≥ *c* for cM threshold *c*, call an IBD segment between sample nodes *i, j*.
4. Stop once there is a single most recent common ancestor.

We simulate the conditional coalescent tree using a discrete time Wright Fisher process with selection. This exact solution as opposed to a coalescent approximation is essential given that those sample nodes with the *de novo* mutation coalescence to a single founder in recent history.

We implement modifications to this algorithm to improve run time. Most sample nodes share very short haplotypes with their common ancestors after tens of generations, so we stop comparing against a node *i* once *r*_*i*_ + *l*_*i*_ < *c*. If the lengths *r*_*i*_ = *r*_*j*_ and *l*_*i*_ = *l*_*j*_ for sample nodes *i, j*, we merge these nodes and compare the merged node against other sample nodes in the preceding steps. This subroutine avoids redundant calculations. We observe in simulations that these implementations lead to approximately linear instead of quadratic compute time. This algorithm makes it feasible to form bootstrap intervals from hundreds of draws for sample size as large as 128,000 diploids.

We use a sample standard deviation ^σ_1:*B*_ from *B* parametric bootstraps to form confidence intervals ŝ ± *q*_α/2_ × ^σ_1:*B*_, where *q*_α/2_ is the normal quantile. The user can specify the number of bootstraps *B* to support the central limit theorem. We do not implement bootstrap bias-correction because the optimization procedure can be unstable for *s* ≤ 0.015. This instability is due to the probability distribution of the IBD rate between *s* ≤ 0.015 model and the neutral *s* = 0.00 model being indistinguishable. This scenario is circumvented by implementing our selection scan beforehand. The apparent normal distribution in Supplementary Figure 12, or a similar probability model with heavier tails, explains why confidence intervals in this our work maintain adequate coverage 1 − α over the selection coefficient *s*.

## Acknowledgements

Research reported in this publication was supported by the National Human Genome Research Institute of the National Institutes of Health under award number HG005701. S.D.T. acknowledges funding support from the National Defense Science and Engineering Graduate Fellowship and National Institute of Health T32 GM081062 Pre-doctoral Training Grant in Statistical Genetics.

The content is solely the responsibility of the authors and does not necessarily represent the official views of the National Institutes of Health. Molecular data for the Trans-Omics in Precision Medicine (TOPMed) program was supported by the National Heart, Lung, and Blood Institute (NHLBI). Core support including centralized genomic-read mapping and genotype calling along with variant quality metrics and filtering were provided by the TOPMed Informatics Research Center (3R01HL-117626-02S1; contract HHSN268201800002I). Core support including phenotype harmonization, data management, sample-identity QC, and general program coordination were provided by the TOPMed Data Coordinating Center (R01HL-120393; U01HL-120393; contract HHSN268201800001I).

We thank Ruoyi Cai, Nobu Masakin, Brian Browning, and Elizabeth Thompson for helpful discussions during the course of this project. We also appreciate Iain Mathieson for feedback on an earlier version of this manuscript.

## Supplemental Acknowledgements

We gratefully acknowledge the studies and participants who provided biological samples and data for the TOPMed project. Funding for the Barbados Asthma Genetics Study was provided by National Institutes of Health (NIH) R01HL104608, R01HL087699, and HL104608 S1. The Mount Sinai BioMe Biobank (BioMe) has been supported by The Andrea and Charles Bronfman Philanthropies and in part by funds from the NHLBI and the National Human Genome Research Institute (NHGRI) (U01HG00638001; U01HG007417; X01HL134588); genome sequencing was funded by contract HHSN268201600037I. The Cleveland Clinic Atrial Fibrillation study (CCAF) was supported by NIH grants R01 HL 090620 and R01 HL 111314, the NIH National Center for Research Resources for Case Western Reserve University and Cleveland Clinic Clinical and Translational Science Award UL1-RR024989, the Cleveland Clinic Department of Cardiovascular Medicine philanthropy research funds, and the Tomsich Atrial Fibrillation Research Fund; genome sequencing was supported by R01HL092577. The Framingham Heart Study (FHS) was supported by contracts NO1-HC-25195, HHSN268201500001I and 75N92019D00031 from the NHLBI and grant supplement R01 HL092577-06S1; genome sequencing was funded by HHSN268201600034I and U54HG003067. The Hypertension Genetic Epidemiology Network Study (Hypergen) is part of the NHLBI Family Blood Pressure Program; collection of the data represented here was supported by grants U01 HL054472, U01 HL054473, U01 HL054495, and U01 HL054509; genome sequencing was funded by R01HL055673. The Jackson Heart Study is supported and conducted in collaboration with Jackson State University (HHSN268201300049C and HHSN268201300050C), Tougaloo College (HHSN268201300048C), and the University of Mississippi Medical Center (HHSN268201300046C and HHSN268201300047C) contracts from NHLBI and the National Institute for Minority Health and Health Disparities (NIMHD); genome sequencing was funded by HHSN268201100037C. The My Life, Our Future samples (MLOF) and data are made possible through the partnership of Bloodworks Northwest, the American Thrombosis and Hemostasis Network, the National Hemophilia Foundation, and Bioverativ; genome sequencing was funded by HHSN268201600033I and HHSN268201500016C. The Venous Thromboembolism project (VTE) was funded in part by grants from the NIH, NHLBI (HL66216 and HL83141) and the NHGRI (HG04735). The Vanderbilt Genetic Basis of Atrial Fibrillation study (VUAF) was supported by grants from the American Heart Association (EIA 0940116N), and grants from the National Institutes of Health (HL092217, U19 HL65962, and UL1 RR024975), and by CTSA award (UL1TR000445) from the National Center for Advancing Translational Sciences; genome sequencing was funded by R01HL092577. The Women’s Health Initiative program (WHI) is funded by NHLBI through contracts 75N92021D00001, 75N92021D00002, 75N92021D00003, 75N92021D00004, 75N92021D00005; genome sequencing was funded by HHSN268201500014C.

## Web Resources

| Name | Description | URL |
| --- | --- | --- |
| CLUES | Method to estimate selection coefficient | https://github.com/standard-aaron/clues |
| hap-ibd | Fast detection of IBD in genetic sequences | https://github.com/browning-lab/hap-ibd |
| ibd-ends | Refines IBD segment endpoints in genetic sequences | https://github.com/browning-lab/ibd-ends |
| IBDkin | Relatedness estimation using IBD segments | https://github.com/YingZhou001/IBDkin |
| IBDNe | Estimation of nonparametric effective sizes using IBD segments | https://faculty.washington.edu/browning/ibdne.html |
| ImaGene | Deep learning for selection | https://github.com/mfumagalli/ImaGene |
| iSAFE | Method to rank SNPs in sweep study | https://github.com/alek0991/iSAFE |
| Relate | Infer ancestral recombination graph | https://myersgroup.github.io/relate/ |
| selscan | Analyzing selection | https://github.com/szpiech/selscan |

The iSWEEP software is freely available under the open source CC0 1.0 Universal License (https://github.com/sdtemple/isweep). This includes a Python package and multiple automated bioinformatics pipelines written in snakemake.

## Author Contributions

S.D.T. planned the study, developed the methods, wrote the software, conducted the analyses, and wrote the manuscript. S.R.B. proposed the study and advised on all aspects of the project, including the methodology and analysis. R.K.W. helped with software development and provided advice for the direction of the study. All authors contributed to editing the manuscript.

---

**Supplementary Figure 1.**
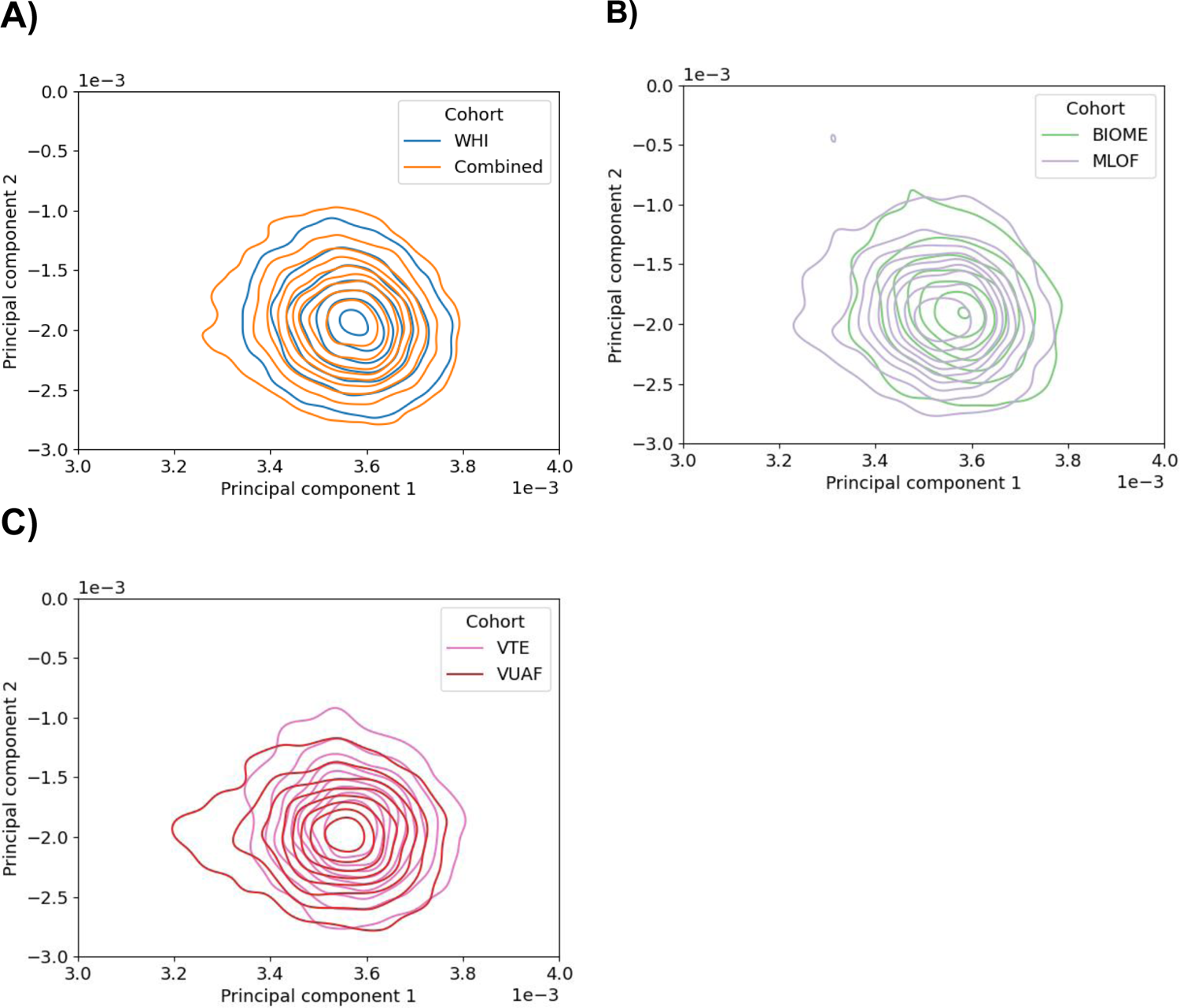
Kernel density plots of principal components with negligible geographic differentiation between European ancestry cohorts Each plot compares two groups: (A) the combined group and WHI; (B) BioMe and MLOF; and (C) VTE and VUAF.

**Supplementary Figure 2.**
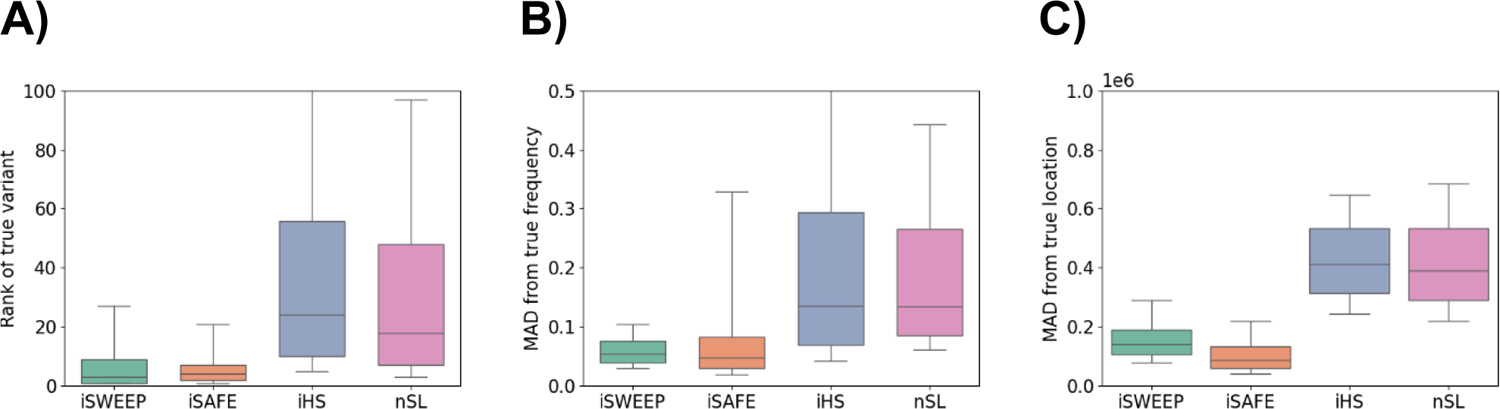
Four methods for identifying the sweeping allele and alleles correlated with it Comparing performance among methods iSWEEP, iSAFE, iHS, and nSL: (A) ranking of the true simulated allele, where 1 is the best rank; (B) mean absolute deviation (MAD) in frequency over the ten best-ranked alleles; and (C) MAD in location over the ten best-ranked alleles. Box plots show 10^th^, 25^th^, 50^th^, 75^th^, and 90^th^ percentiles from one thousand simulations of population bottleneck effective sizes and selection coefficient *s* ∈ [0.015,0.020,0.025,0.030,0.035]. The sample size is five thousand diploids. Population bottleneck (BN) is the demographic scenario. Mutation, recombination, and gene conversion rates are 1e-8, 1e-8, and 2e-8.

**Supplementary Figure 3.**
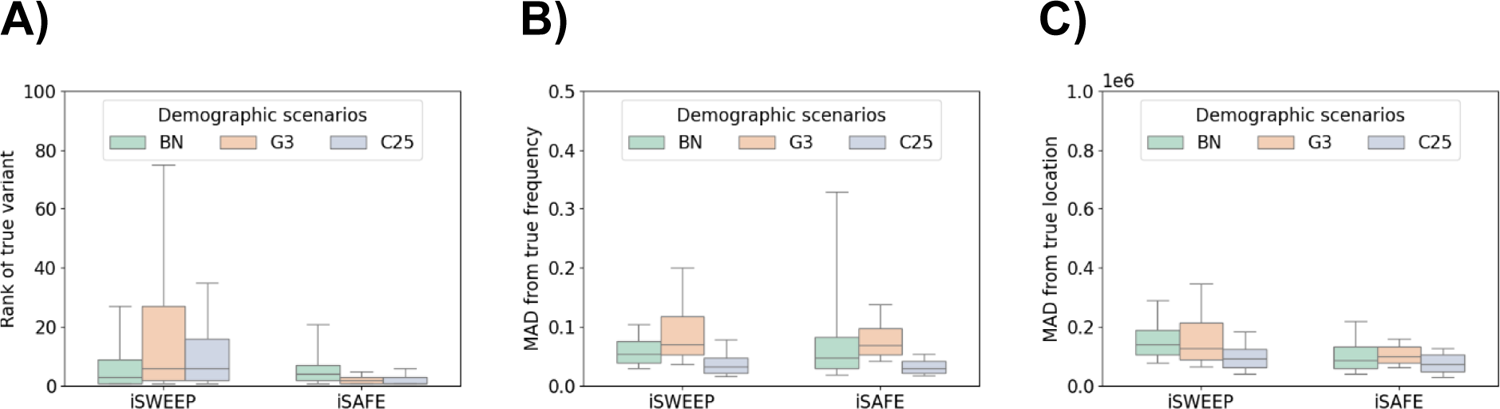
Identifying the sweeping allele in different demographic scenarios Comparing performance among methods iSWEEP and iSAFE: (A) ranking of the true simulated allele, where 1 is the best rank; (B) mean absolute deviation (MAD) in frequency over ten best-ranked alleles; and (C) MAD in location over ten best-ranked alleles. Box plots show 10^th^, 25^th^, 50^th^, 75^th^, and 90^th^ percentiles from two hundred simulations of selection coefficient *s* = 0.03 and different demographic scenarios: population bottleneck (BN), three phases of exponential growth (G3), constant size *N*_*e*_ = 25,000 (C25). The sample size is five thousand diploids. Mutation, recombination, and gene conversion rates are 1e-8, 1e-8, and 2e-8.

**Supplementary Figure 4.**
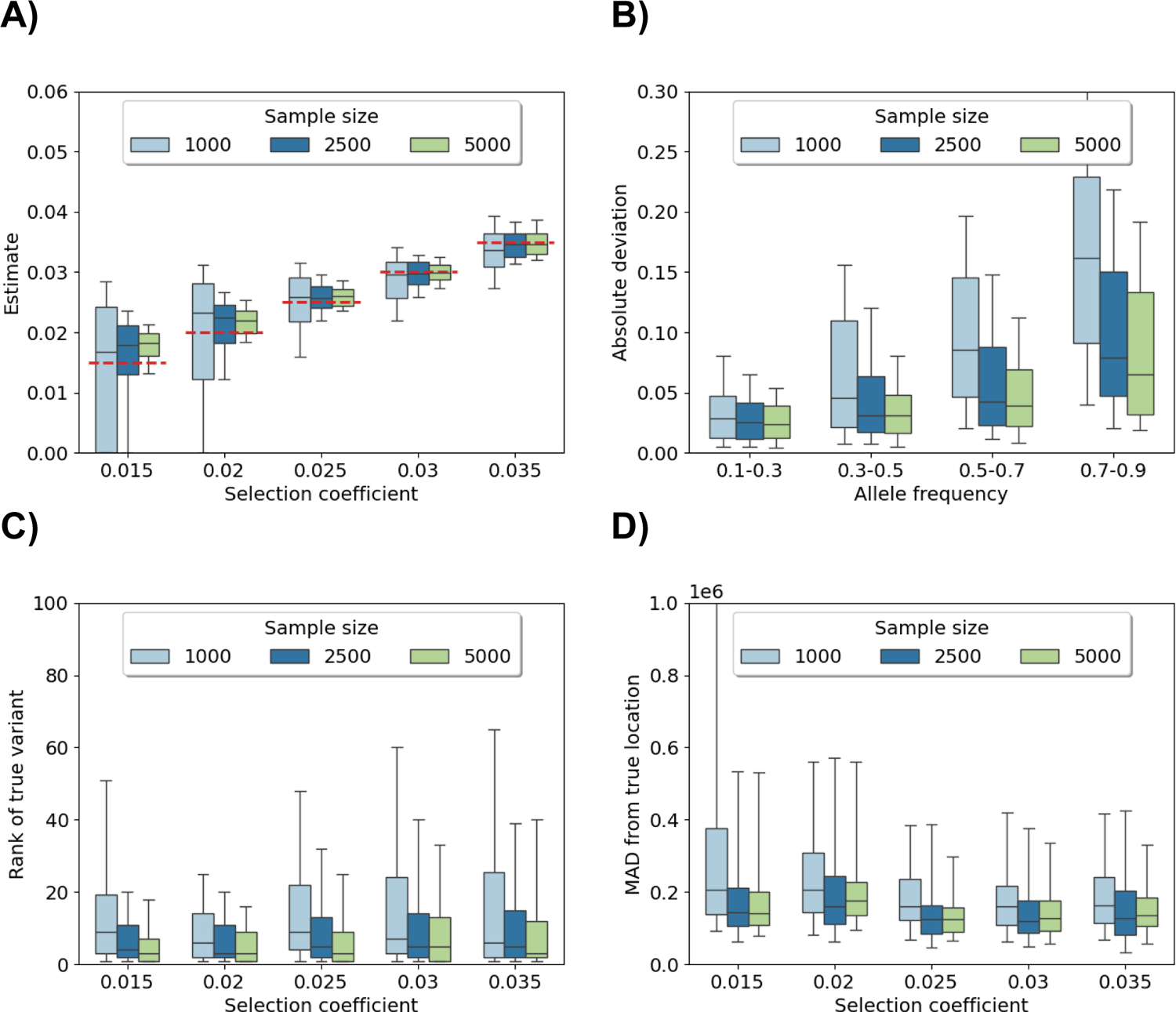
Performance of **iSWEEP** for increasing sample size Comparing between sample sizes 1000 (light blue), 2500 (dark blue), and 5000 (light green) diploids: (A) estimating selection coefficients, (B) estimating the frequency of the sweeping allele, (C) ranking the true simulated allele, and (D) mean absolute deviation (MAD) in location among ten best-ranked alleles. Box plots show 10^th^, 25^th^, 50^th^, 75^th^, and 90^th^ percentiles from two hundred simulations of each selection coefficient. Horizontal dashed red lines correspond to the true selection coefficient. Population bottleneck (BN) is the demographic scenario. Mutation, recombination, and gene conversion rates are 1e-8, 1e-8, and 2e-8.

**Supplementary Figure 5.**
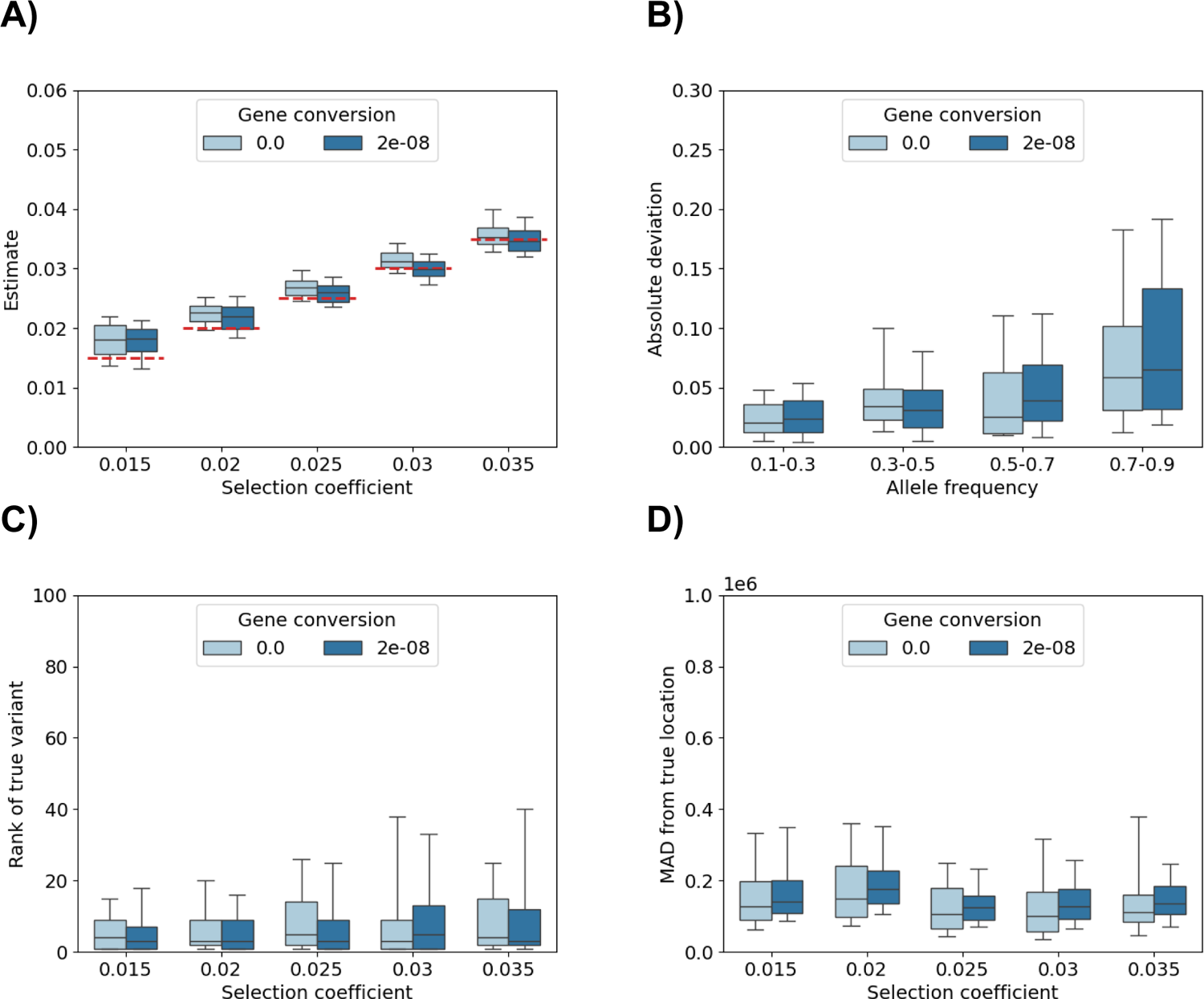
Performance of **iSWEEP** with and without gene conversion Comparing gene conversion rates of zero (light blue) versus 2e-8 (dark blue): (A) estimating selection coefficients, (B) estimating the frequency of the sweeping allele, (C) ranking the true simulated allele, and (D) mean absolute deviation (MAD) in location among lowest ten sorted alleles. Box plots show 10^th^, 25^th^, 50^th^, 75^th^, and 90^th^ percentiles from sixty simulations of each selection coefficient. Horizontal dashed red lines correspond to the true selection coefficient. Sample size is five thousand diploids. Population bottleneck (BN) is the demographic scenario. Mutation and recombination rates are 1e-8 and 1e-8.

**Supplementary Figure 6.**
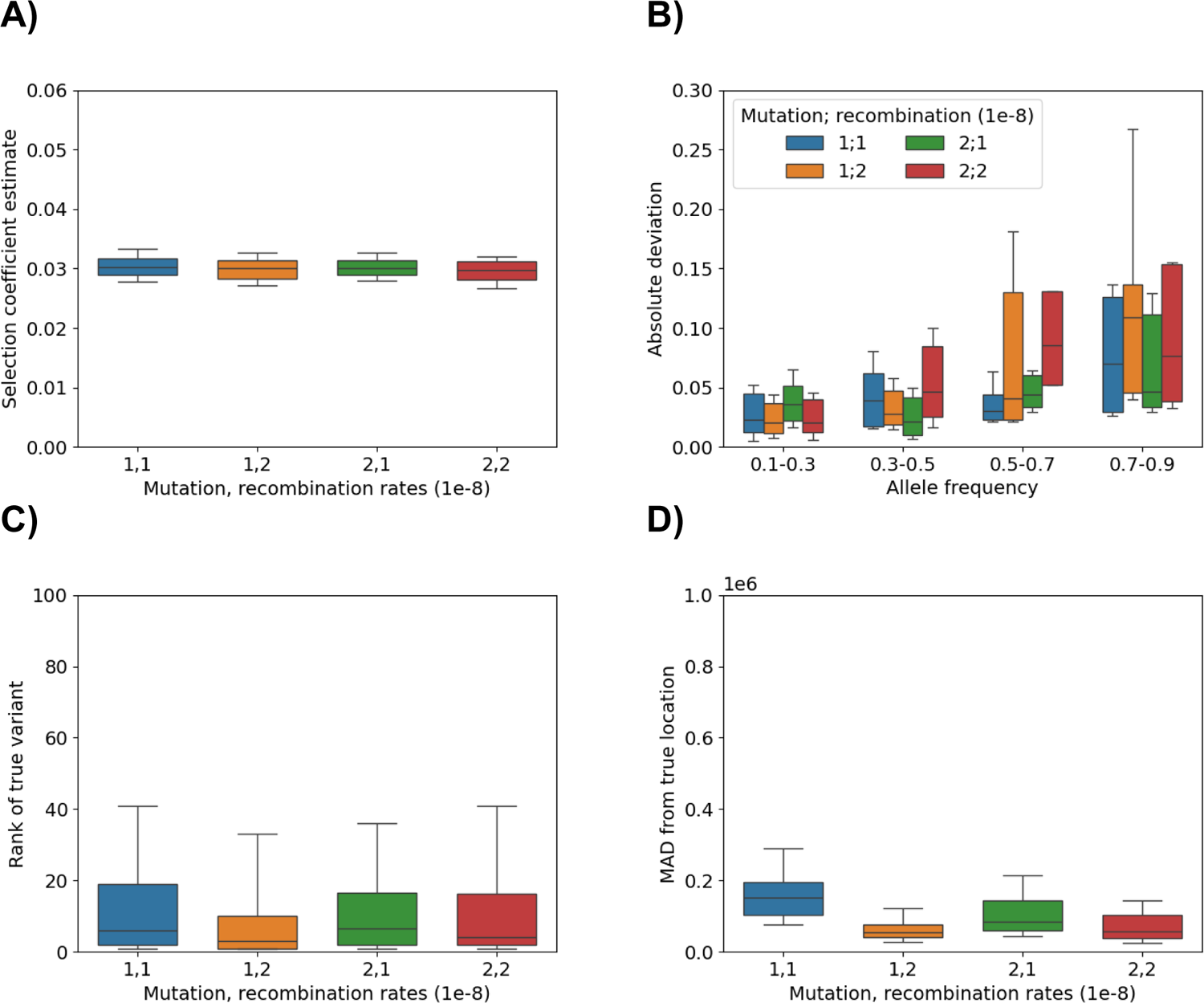
Performance of **iSWEEP** for different mutation and recombination rates Comparing pair combinations of mutation rate μ and recombination rate ρ: (A) estimating the selection coefficient *s* = 0.03, (B) estimating the frequency of the sweeping allele, (C) ranking the true simulated allele, and (D) mean absolute deviation (MAD) in location among lowest ten sorted alleles. Blue is rates (μ, ρ) = (1e-8,1e-8), orange is rates (1e-8,2e-8), green is rates (2e-8,1e-8), and red is rates (2e-8,2e-8). There are sixty simulations for each setting. Violin plot in (A) uses default settings in the seaborn Python package. Box plots in (B-D) show 10^th^, 25^th^, 50^th^, 75^th^, and 90^th^ percentiles. Sample size is five thousand diploids. Sample size is five thousand diploids. Population bottleneck (BN) is the demographic scenario. Gene conversion rate is 2e-8.The selection coefficient is *s* = 0.03.

**Supplementary Figure 7.**
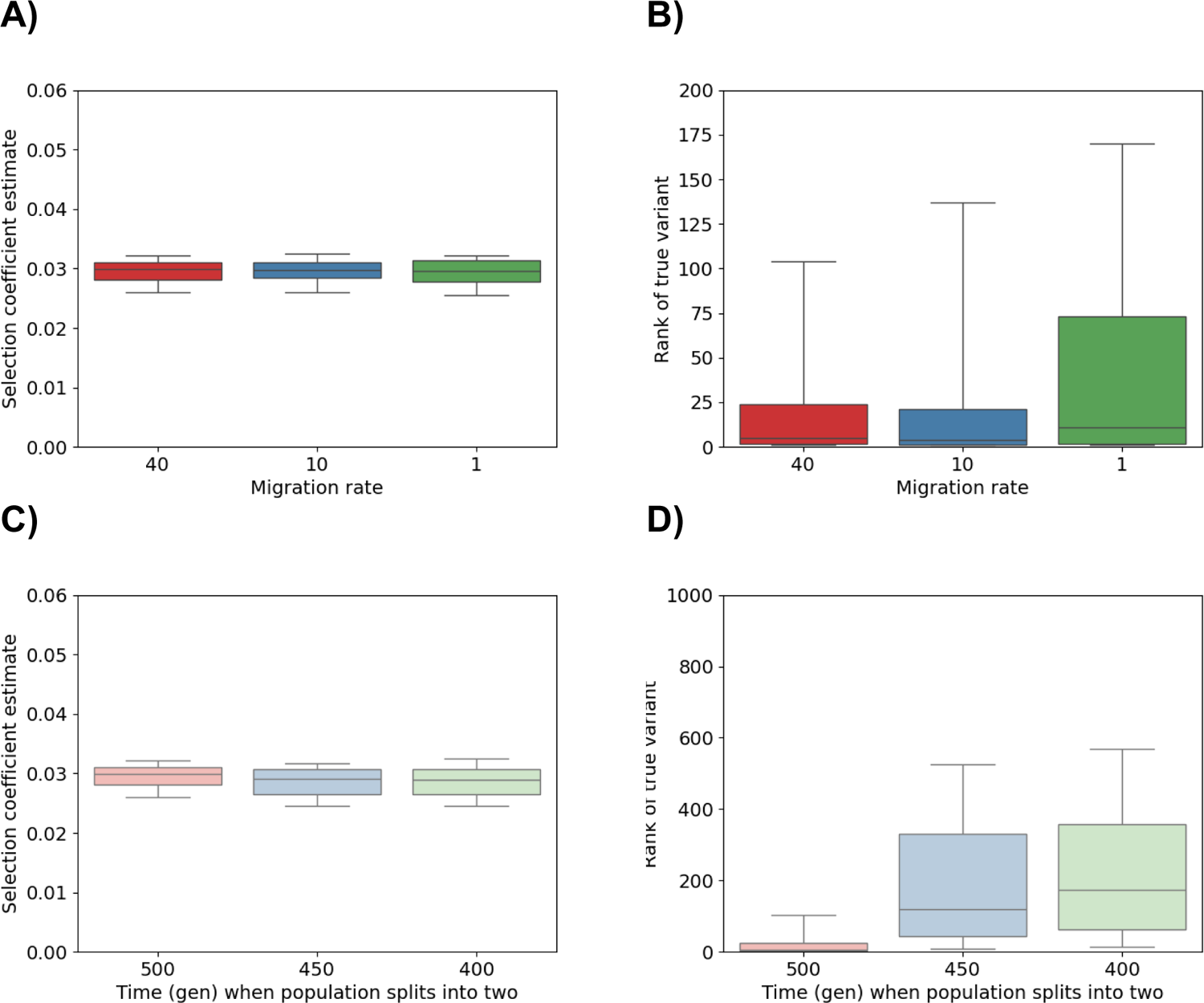
Performance of **iSWEEP** for different scenarios of cryptic population substructure Evaluating accuracy at estimating the selection coefficient *s* = 0.03 (A,C) and ranking the true simulated allele (B,D) when two subpopulations split 500 generations ago and migrate between each other at the rates forty, ten, and one percent (A,B) or when two subpopulations split 500, 450, and 400 generations ago and migrate between each other at the rate forty percent (C,D). There are two hundred simulations for each setting. The box plot in (B) shows 10^th^, 25^th^, 50^th^, 75^th^, and 90^th^ percentiles. Sample size is five thousand diploids. Population bottleneck (BN) is the demographic scenario. Mutation, recombination, and gene conversion rates are 1e-8, 1e-8, and 2e-8. Note that the *y*-axis scale is changed in (B) and (D).

**Supplementary Figure 8.**
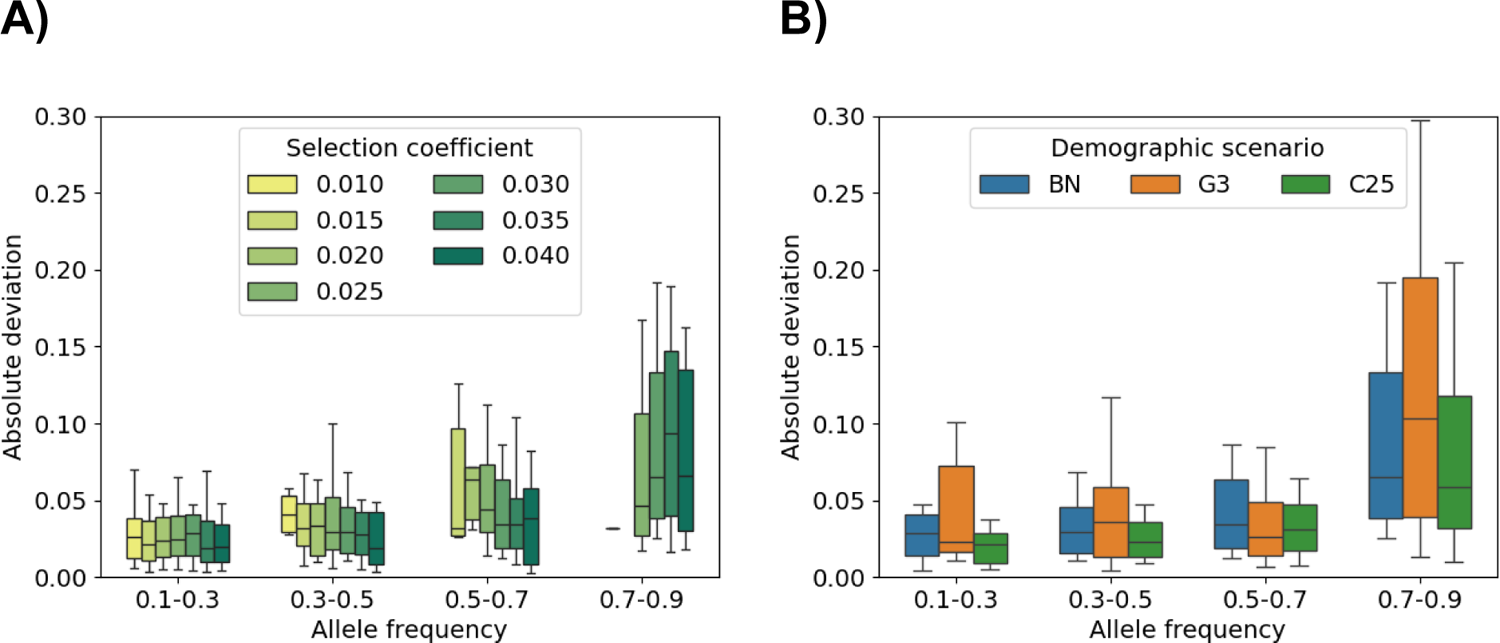
Estimating the frequency of the unknown sweeping allele Absolute deviations between allele frequency estimate and the truth. (A) Results aggregated over two hundred simulations each for *s* ∈ [0.01,0.015,0.02,0.025,0.03,0.035,0.04] and demographic scenarios. (B) Results aggregated over two hundred simulations each for fixed *s* = 0.03 and demographic scenarios population bottleneck (BN), three phases of exponential growth (3G), constant size *N*_*e*_ = 25,000, and two subpopulations with forty percent migration (M40). Box plots show 10^th^, 25^th^, 50^th^, 75^th^, and 90^th^ percentiles. Sample size is five thousand diploids. Mutation, recombination, and gene conversion rates are 1e-8, 1e-8, and 2e-8.

**Supplementary Figure 9.**
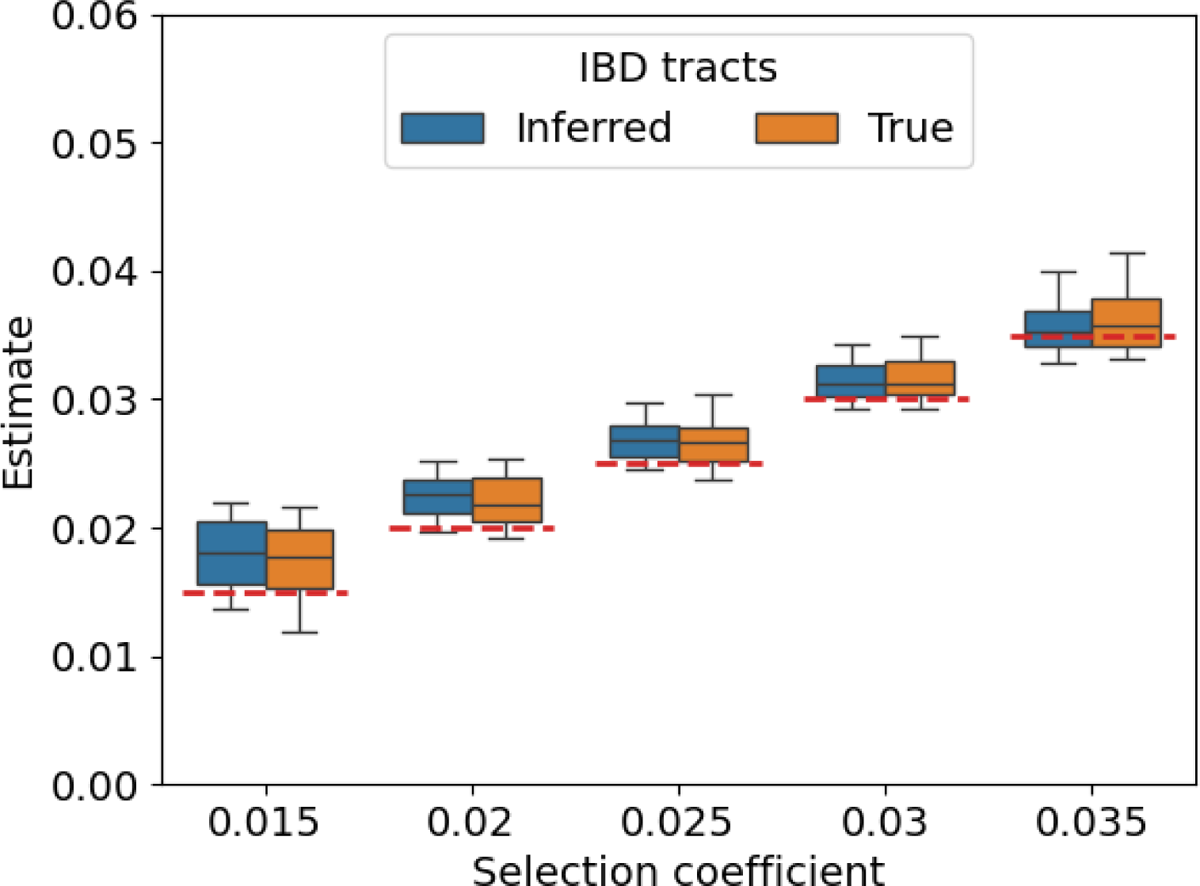
Estimating selection coefficients using true versus inferred IBD Results from iSWEEP based on simulations of zero gene conversion and either inferred IBD tracts from hap-ibd and ibd-ends (blue) or true IBD tracts from tskibd (orange). Box plots show 10^th^, 25^th^, 50^th^, 75^th^, and 90^th^ percentiles based on sixty simulations for each selection coefficient. Horizontal dashed red lines correspond to the true selection coefficient. Sample size is five thousand diploids. Population bottleneck (BN) is the demographic scenario. Mutation and recombination rates are 1e-8 and 1e-8.

**Supplementary Figure 10.**
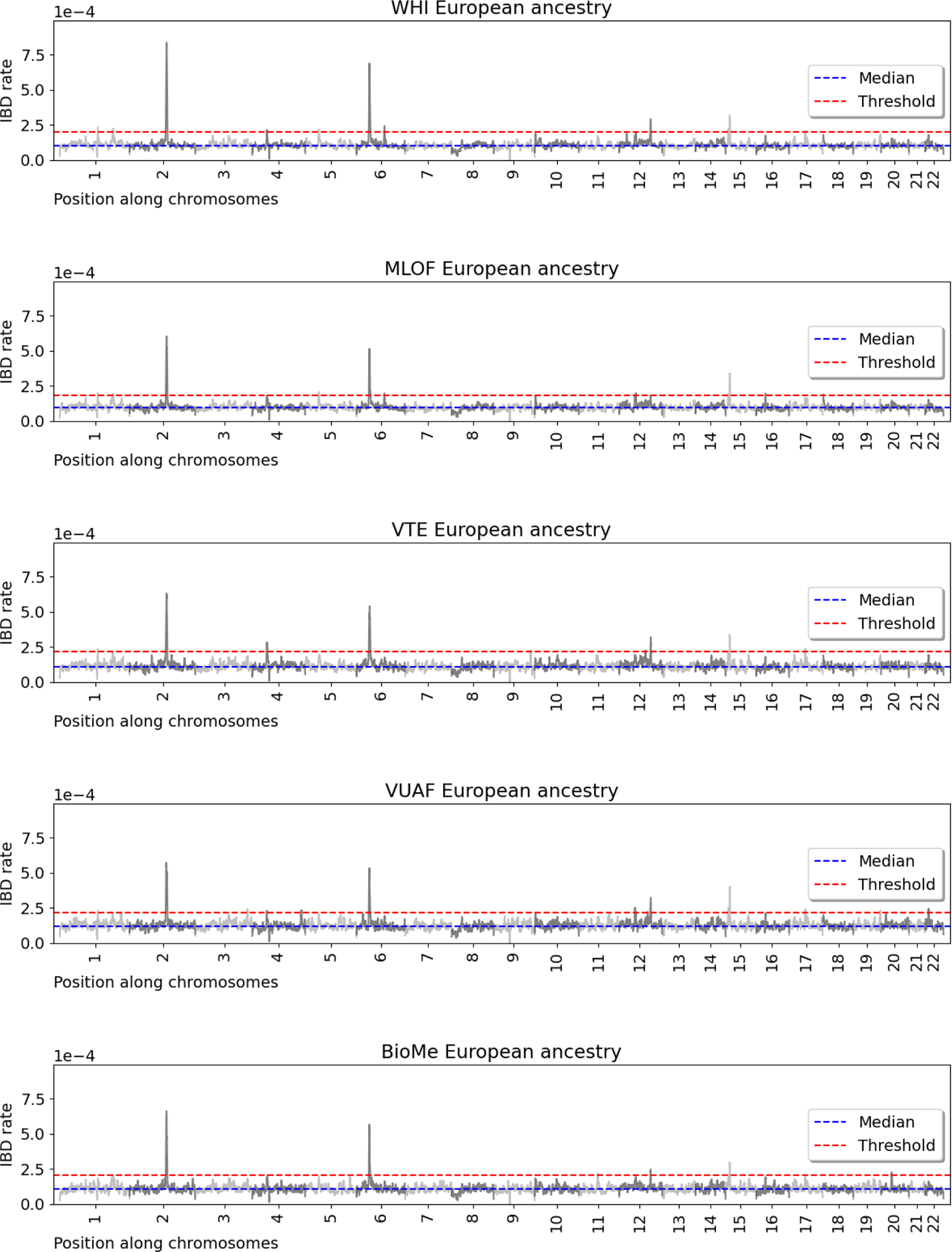
Cohort study selection scans of TOPMed European Americans

**Supplementary Figure 11.**
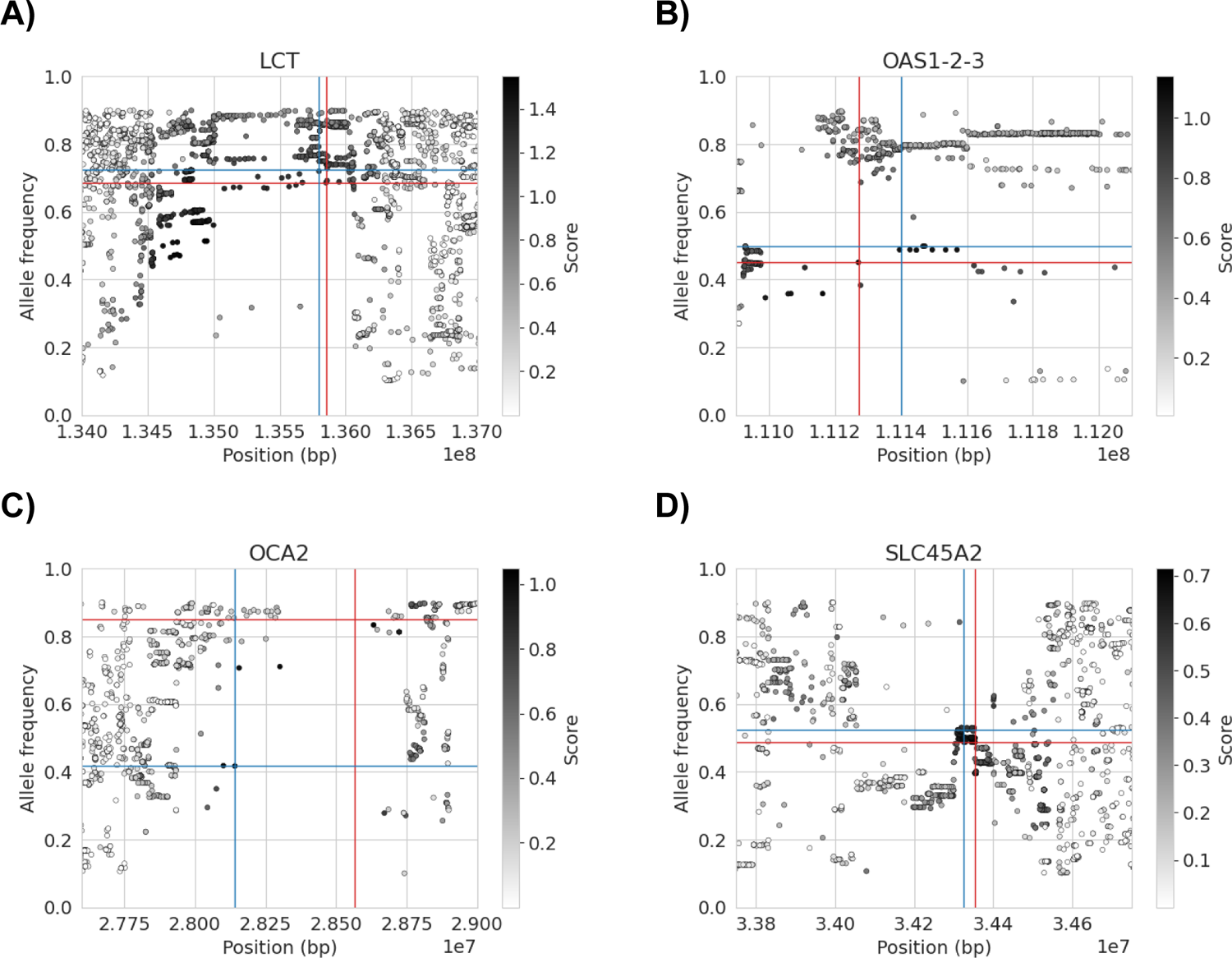
Spectrum of common variants at selected loci Position in basepairs (*x*-axis) by allele frequency (*y*-axis) for common variants at loci passing the selection scan: (A) LCT; (B) OAS1-2-3; (C) OCA2; and (D) SLC45A2. Color bars show the *Z*_*j*_ scores, where darker grays correspond to greater scores. Blue lines denote the haplotype-based allele frequency and location estimates. Red lines denote frequency and location of the SNP with the greatest *Z*_*j*_ score. Results shown for combined EUR group (A, B, D) and for WHI cohort (C).

**Supplementary Figure 12.**
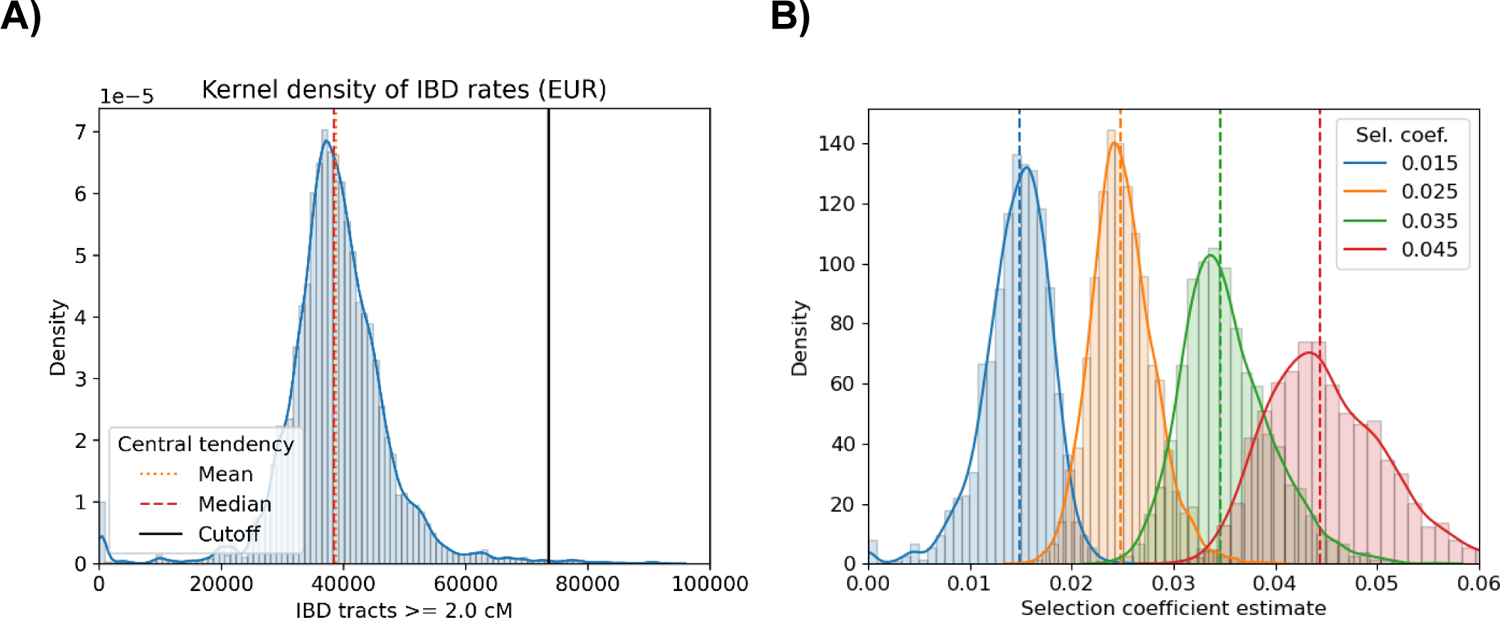
Identity-by-descent rates and selection coefficient estimates look to be normally distributed (A) Histogram and fitted curve for inferred IBD rates along the genome in TOPMed EUR ancestry samples are shown. (B) Histograms and fitted curve for selection coefficient estimates in simulations using EUR estimated effective sizes and *p*(0) = 0.50 are shown.

**Supplementary Figure 13.**
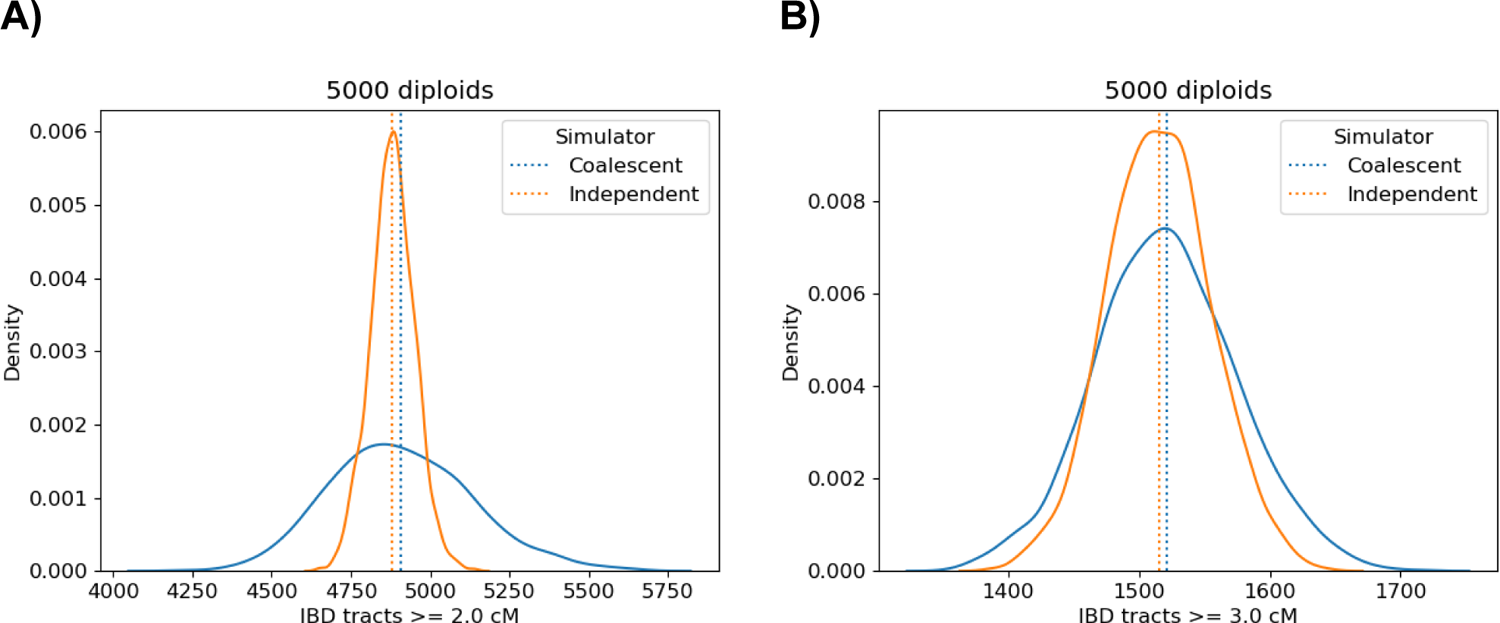
The parametric bootstrap reflects the true variance in the identity-by-descent process Kernel density estimates of the number of IBD tracts ≥ 2.0 and ≥ 3.0 cM (A and B) over 2,500 replicate simulations of five thousand diploids in the BN demographic scenario are shown. Results from the coalescent simulator for IBD tracts (blue) and a simulator for independent IBD tracts (orange) are shown. Vertical dashed lines denote means.

**Supplementary Table 1.** Algorithm settings for detecting IBD segments in sequence data. Unless otherwise specified in this table, we use the defaults settings in hap-ibd and ibd-ends. We use hap-ibd version 1.0 from May 10, 2023 and ibd-ends version 1.1 from May 20, 2022. The IBD segments ≥ 2.0 and ≥ 3.0 cM are a filtered subset of the ibd-ends output. The error rate in ibd-ends should be modified if in a preliminary data analysis the estimated error rate differs from the default by more than a factor of three.

| cM segment type | Software | Algorithm settings | Purpose |
| --- | --- | --- | --- |
| $\geq 2.0$ and $\geq 3.0$ cM | hap-ibd | min-seed=0.5<br>min-extend=0.2<br>min-output=1.0<br>min-maf=0.4 | Candidate segments for ibd-ends |
| $\geq 2.0$ and $\geq 3.0$ cM | ibd-ends | min-maf=0.001<br>err=2e-4<br>Median endpoints | Selection scan, estimation |
| $\geq 1.0$ cM | hap-ibd | min-seed=0.5<br>min-extend=0.2<br>min-output=1.0<br>min-maf=0.1 | IBD outgroup algorithm |

**Supplementary Table 2.**
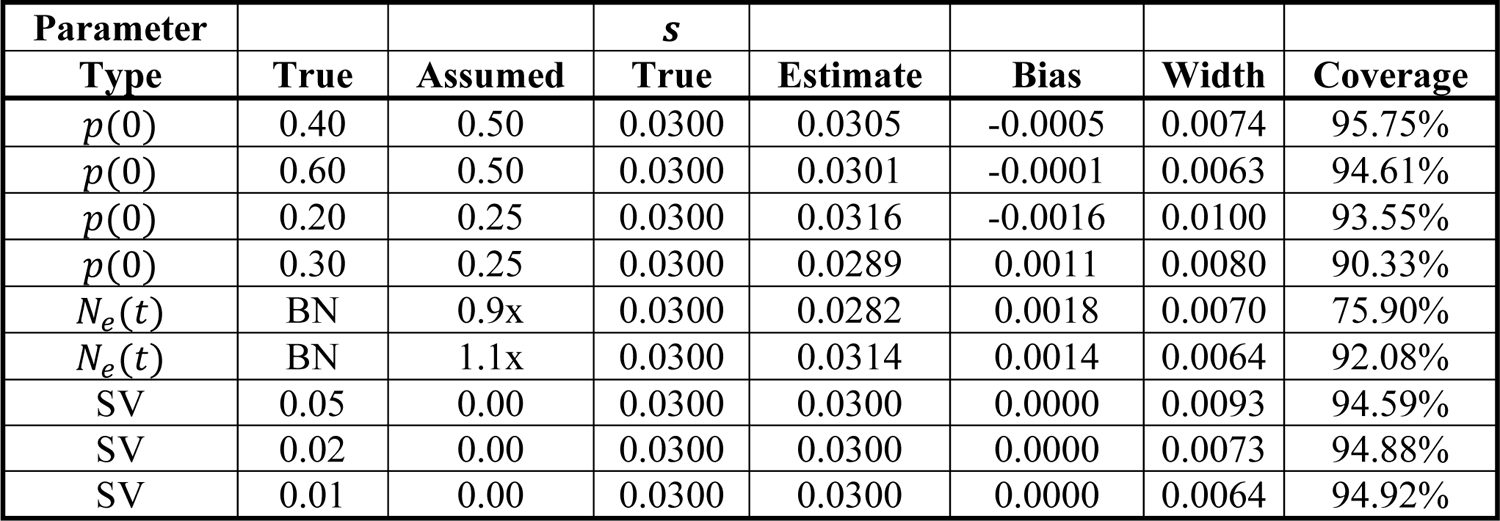
Selection coefficient estimation based on true IBD segments and model misspecification Each row represents the results of 10,000 replicates. Summary statistics are the mean estimate, the mean bias, the width of confidence interval, and the percentage of intervals containing the true selection coefficient. Ninety-five percent confidence intervals are based on fifty bootstraps. Default settings are *s* = 0.03, *p*(0) = 0.50, population bottleneck demographic history (BN), the multiplicative genetic model, and an ongoing sweep from a new beneficial mutation. The sample size is five thousand diploids. The Assumed column is the parameter value used in conditional estimation of *s*. Abbreviations 0.9x and 1.1x indicate that the *N*_*e*_(*t*) is uniformly scaled by that value. SV concerns the allele frequency of standing neutral variation.

**Supplementary Table 3.** Selection coefficient estimation based on IBD segments inferred from sequence data and different demographic scenarios For each row, estimates Ŝ and coverage are averaged over one hundred simulations. Ninety-five percent confidence intervals are based on fifty bootstraps. Time refers to generations ago when the *de novo* mutation arose. Demographic scenarios in the first column abbreviated as bottleneck (BN), three phases of exponential growth (G3), constant size *N*_*e*_ = 25,000 (C25), and random mixing with proportion *m* = 0.40 between two subpopulations. The sample size is five thousand diploids. Mutation, recombination, and gene conversion rates are 1e-8, 1e-8, and 2e-8.

| $N_e(t)$ | Time | True $s$ | Estimate $\bar{\hat{s}}$ | Coverage |
| --- | --- | --- | --- | --- |
| BN | 250 | 0.030 | 0.0304 | 98 |
| BN | 300 | 0.030 | 0.0295 | 86 |
| C25 | 250 | 0.030 | 0.0311 | 97 |
| C25 | 300 | 0.030 | 0.0300 | 95 |
| G3 | 250 | 0.030 | 0.0292 | 91 |
| G3 | 300 | 0.030 | 0.0279 | 66 |
| $m = 0.4$ | 250 | 0.030 | 0.0294 | 79 |
| $m = 0.4$ | 300 | 0.030 | 0.0291 | 87 |

**Supplementary Table 4.** Algorithm settings for simulation study on sequence data Average CPU time for iHS, iSAFE, iSWEEP, and nSL is based on four *s* = 0.03 simulations of 5000 samples. (Wall clock time for iSWEEP is faster if running hap-ibd in parallel across cores.) CLUES and ImaGene are analyzed using 500 of 5000 samples due to runtime limitations. All methods except ImaGene require a genetic map; ImaGene requires a genome-wide recombination rate.

| Name | Purpose | Approach | Input data | Assumptions | CPU time |
| --- | --- | --- | --- | --- | --- |
| iHS | Ranking SNPs | EHH-based | Phased variants | Derived allele known | 107.01 min |
| iSAFE | Ranking SNPs | EHH-based | Phased variants | Derived allele known | 34.07 min |
| iSWEEP | Ranking SNPs | IBD-based | Phased variants;<br>$\geq 1.0$ cM IBD segments | Inferred adaptive allele | 5.83 min |
| nSL | Ranking SNPs | EHH-based | Phased variants | Derived allele known | 22.00 min |
| iSWEEP | Estimating $p(0)$ | IBD-based | Phased variants;<br>$\geq 1.0$ cM IBD segments | Inferred adaptive allele | 11.75 sec |
| CLUES | Estimating $s$ | Modeling coalescent | Inferred ARGs | Causal allele known | More than half a day |
| ImaGene | Estimating $s$ | Deep learning | Simulated training data | N/A | $\geq 1$ week training the network |
| iSWEEP | Estimating $s$ | Modeling IBD segments | Phased variants;<br>$\geq 3.0$ cM IBD segments | Estimates for $p(0)$ and focal location | 2.79 min |

**Supplementary Table 5.**
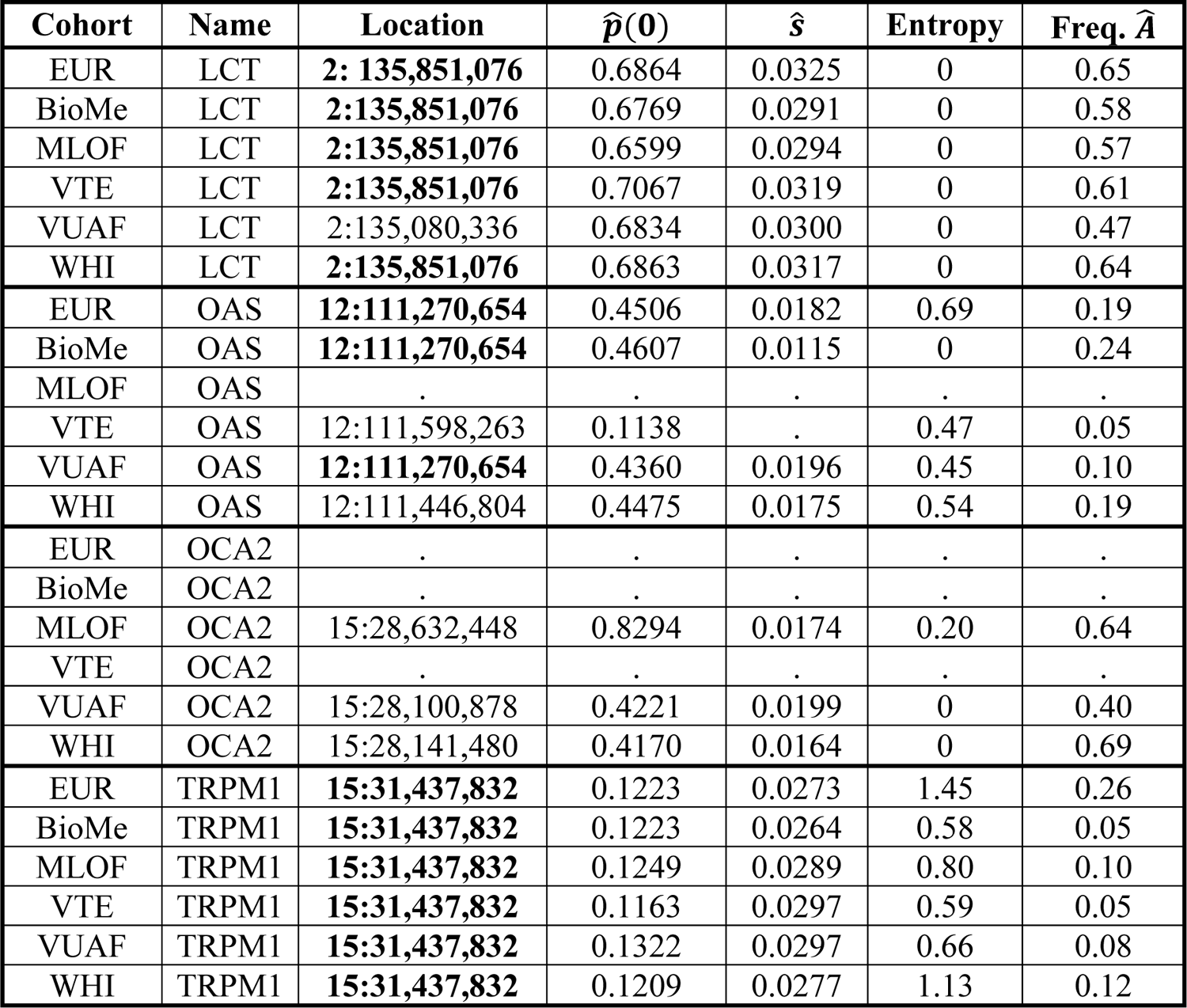
Cohort-specific estimation for TOPMed European Americans IBD entropy, the proportion of haplotypes in the inferred IBD outgroup Â, and allele frequency, location, and selection coefficient estimates for selected loci LCT, OAS, OCA2, and TRPM1. Either IBD entropy is less than 0.50 *or* there is consensus across studies of the best-ranked variant for these loci. Boldface marks SNPs that are the best-ranked among multiple cohort-specific analyses. N/A marks if the locus does not pass the selection scan in an analysis.

